# Spectral partitioning identifies individual heterogeneity in the functional network topography of ventral and anterior medial prefrontal cortex

**DOI:** 10.1101/651117

**Authors:** Claudio Toro-Serey, Sean M. Tobyne, Joseph T. McGuire

## Abstract

Regions of human medial prefrontal cortex (mPFC) and posterior cingulate cortex (PCC) are part of the default network (DN), and additionally are implicated in diverse cognitive functions ranging from autobiographical memory to subjective valuation. Our ability to interpret the apparent co-localization of task-related effects with DN-regions is constrained by a limited understanding of the individual-level heterogeneity in mPFC/PCC functional organization. Here we used cortical surface-based meta-analysis to identify a parcel in human PCC that was more strongly associated with the DN than with valuation effects. We then used resting-state fMRI data and a data-driven network analysis algorithm, spectral partitioning, to partition mPFC and PCC into “DN” and “non-DN” subdivisions in individual participants (n = 100 from the Human Connectome Project). The spectral partitioning algorithm identified individual-level cortical subdivisions that varied markedly across individuals, especially in mPFC, and were reliable across test/retest datasets. Our results point toward new strategies for assessing whether distinct cognitive functions engage common or distinct mPFC subregions at the individual level.

**Highlights:**

- The topography of Default Network cortical regions varies across individuals.
- A community detection algorithm, spectral partitioning, was applied to rs-fMRI data.
- The algorithm identified individualized Default Network regions in mPFC and PCC.
- Default Network topography varied across individuals in mPFC, moreso than in PCC.
- Overlap of task effects with DN regions should be assessed at the individual level.

## 1. Introduction

Human medial prefrontal cortex (mPFC) and posterior cingulate cortex (PCC) are jointly associated with a diverse set of cognitive processes (Hiser & Koenigs, 2018; Kragel et al., 2018), and contain subregions that are part of the brain’s default network (Buckner & DiNicola, 2019; DN; Buckner et al., 2008). DN regions are characterized by a decrease in BOLD activity during externally oriented tasks that require attention or cognitive control, in comparison with less-demanding task conditions or periods of rest (Buckner et al., 2008; Laird et al., 2009; McKiernan et al., 2003). The DN can also be identified on the basis of a distinctive pattern of inter-region correlations in resting-state fMRI data (Buckner et al., 2008; Fox et al., 2005; Greicius et al., 2003; Yeo et al., 2011).

Many different cognitive task manipulations evoke patterns of brain activity that overlap with DN regions in ventral and anterior mPFC and in PCC. Examples include manipulations of self-referential thinking (Gusnard et al., 2001; Mitchell et al., 2005), memory (Euston et al., 2012; Schacter et al., 2007), affective regulation (Reddan et al., 2018; Schiller et al., 2008), and subjective valuation (Bartra et al., 2013; Clithero & Rangel, 2014; Kable & Glimcher, 2007; Levy et al., 2011). Some of these task-related effects are thought to reflect processes integral to the functional role of the DN, such as internally oriented cognition, scene construction, and self-projection (Buckner & Carroll, 2007; Hassabis & Maguire, 2007). For other task-related effects, such as subjective valuation (i.e., greater BOLD activity in response to more highly valued choice prospects and outcomes, relative to prospects and outcomes that are less highly valued), the degree of overlap with DN regions is only partial and the reason for the overlap is less obvious (Acikalin et al., 2017). Insofar as the DN shares a subset of nodes in common with the distributed brain systems that support valuation and other functions, this has potential to inform our theoretical understanding of the cognitive operations involved in those functions (Northoff & Hayes, 2011). As a result, there is widespread interest in understanding the degree to which DN regions overlap topographically with task-related effects (Buckner & DiNicola, 2019; DiNicola et al., 2019; Spreng, 2012).

However, strong conclusions about functional colocalization require consideration of individual-level heterogeneity in topographic patterns of brain activity. A recognized limitation of group averaging and meta-analysis is that the functional topography of individual brains can be misaligned and blurred (Fedorenko et al., 2012; Guntupalli et al., 2018; Michalka et al., 2015; Tobyne et al., 2018; Wang et al., 2015; Woo et al., 2014), exaggerating the apparent overlap across domains. This concern is especially pronounced in ventral mPFC, which is subject to considerable idiosyncratic cortical folding (Lopez-Persem et al., 2019; Mackey & Petrides, 2014; Zilles et al., 2013) and inter-subject functional variability (Mueller et al., 2013). An alternative approach is to focus on analyses at the individual-participant level. Individual-level analyses of fMRI data have identified idiosyncratic, reliable, and valid patterns of functional organization that would be blurred in aggregative estimates (Gordon et al., 2017; Gratton et al., 2018; Laumann et al., 2015; Michalka et al., 2015; Tobyne et al., 2018), and subject-specific network arrangements have been found to predict behavioral characteristics (Kong et al., 2018). Recent work has uncovered fine-grained subdivisions within the DN using both data-driven clustering and individually customized seed-based connectivity analysis (Braga & Buckner, 2017; Braga et al., 2019). It is therefore possible that the apparent overlap of the DN with task-related effects might, in some cases, be attributable to low effective spatial resolution, and that the organization of mPFC and PCC might be better understood at the individual level. An important first step in investigating this possibility, and the goal of the present paper, is to quantify the degree of variability in the topography of the DN within mPFC and PCC across a large sample of individuals.

A useful way to characterize individual-specific brain organization is to examine patterns of resting-state functional connectivity. Connectome-based analyses of resting-state functional connectivity have been fruitful in identifying individualized functional subregions that correspond well to task-induced activity patterns (Gordon et al., 2017; Laumann et al., 2015; Smith et al., 2009; Tobyne et al., 2018). A functional connectome can be represented in the form of a network, and graph theoretic methods can be applied to analyze the network’s structure (Bassett & Sporns, 2017; Rubinov & Sporns, 2010). In the context of network analysis, community detection algorithms subdivide brain networks into sets of nodes that share more connections with each other than with the rest of the network (Fortunato & Hric, 2016; Garcia et al., 2018). Here we use the technique of spectral partitioning (SP), an efficient community detection algorithm that deterministically subdivides a network into two communities (Belkin & Niyogi, 2003; Chung, 1997; Fiedler, 1975). SP has previously been used to characterize the posterior-anterior functional gradient of the insula using resting-state fMRI data (Tian & Zalesky, 2018), and was shown to robustly and reliably separate both simulated and actual primate ECoG networks (Toker & Sommer, 2019). We use SP here to identify subsets of nodes within mPFC and PCC that share spontaneously covarying temporal activation patterns during rest.

In this study, we aimed to subdivide mPFC and PCC into individual-specific DN and non-DN communities, and to quantify the degree of topographic heterogeneity in the resulting community structure over time and across individuals. We did this by capitalizing on the respective strengths of meta-analysis and subject-specific analyses of brain networks. We used a data-driven network-analysis procedure to identify two communities that each spanned both mPFC and PCC in each individual participant. We found that the resulting communities had a stereotyped topographic layout within PCC (according to a label-agnostic similarity metric), whereas their layout in mPFC was variable across individuals but stable across test/re-test. We took advantage of the more consistent configuration within PCC to assign meta-analysis-derived labels to the two communities. Because our data-driven method established correspondence between PCC subregions and mPFC subregions, the labels defined in PCC could then be indexed into the more heterogeneous community structure of mPFC in each individual.

The outline of our paper is as follows. First, we defined a search space by selecting parcels from an established brain atlas (Glasser et al., 2016) that corresponded to previously defined DN and limbic networks on the medial cortical wall (Yeo et al., 2011). A cortical surface-based meta-analysis of the DN and valuation literatures identified a parcel in PCC that was DN-specific at the aggregate level. Valuation was selected as an example of a cognitive domain in which group-average activity patterns overlap extensively with DN regions on the medial surface, despite being segregable elsewhere (Acikalin et al., 2017). We then derived a functional connectivity network of all the surface vertices within the search space for each of 100 individual resting-state fMRI data sets from the Human Connectome Project (HCP; Van Essen et al., 2012), and used the SP algorithm to subdivide each individual’s network into DN and non-DN communities (labeled according to which community included the meta-analytically identified DN-specific parcel in PCC). Focusing on individual vertices in the search space rather than the parcels (as is typical in brain network analyses) allowed us to finely delineate the topographic extent of each community. The resulting communities varied topographically across individuals, while also appearing to follow common organizational principles. Test-retest analyses showed that these partitionings were similar across scanning days within (but not between) individuals, and that individual-level idiosyncrasy was greater in mPFC. Partitionings obtained from the SP algorithm had higher test-retest reliability than did analogous results from seed-based functional connectivity. We observed a trend for the DN community to be located within principal sulci in ventral mPFC and left PCC, but in gyri within superior mPFC and right PCC. Lastly, we describe how the structure of the resulting automatically defined DN and non-DN communities both aligns with and differs from a recently proposed scheme for identifying subdivisions within the DN (Braga & Buckner, 2017; Braga et al., 2019). Our work highlights the usefulness of estimating brain effects at the individual level in mPFC and PCC, and provides a new framework and tool set for future investigations of overlap across cognitive domains.

## 2. Material and Methods

### 2.1. Data and Code Accessibility Statement

All code used in this study is openly available at https://github.com/ctoroserey/mPFC_partitioning. Resting-state fMRI data were obtained from the Human Connectome Project (Van Essen et al., 2012).

### 2.2. Search space

For all analyses, we defined our search space based on the 17-network parcellation proposed by Yeo et al. (2011). First, we selected vertices on the medial cortical surface that were contained by the DN and limbic networks in HCP’s 32,000 vertex surface space (fs_LR_32k). Next, we overlaid those networks on a parcellated atlas of the human cortical surface (360 regions; Glasser et al., 2016), and retained a set of parcels that covered approximately the same brain regions (visually inspected, retaining parcels that appeared to have at least 15% overlap). This resulted in a search space that consisted of 40 parcels across hemispheres (Supplementary Table 1). The search space in each hemisphere was naturally divided into two spatially non-contiguous clusters in PCC and mPFC, facilitating the examination of each region separately.

### 2.3. Meta-analysis

We used a novel approach to cortical surface parcel-based meta-analysis to assess whether individual parcels within the search space were preferentially associated with subjective valuation or with decreased activity during externally oriented tasks, which served to operationalize the DN. For subjective valuation, we gathered peak activation coordinates from 200 studies that reported positive effects in contrasts of higher-value minus lower-value outcomes or prospects (Bartra et al., 2013). For the DN, we acquired coordinates from 80 studies that reported reductions in BOLD during externally directed tasks compared to a baseline (Laird et al., 2009). The coordinates represent areas that exceeded the statistical significance threshold in each original study. For each study, we created an indicator map in standard volumetric space (MNI152, 1 mm resolution) which contained values of 1 in a 10 mm radius sphere around each reported activation peak, and values of 0 elsewhere (Wager et al., 2009). The indicator map for each study was then projected to a standard cortical mesh (fsaverage, 160,000 vertices, projfrac-max from 0 to 1 by 0.25, registered using mni152.register.dat) using FreeSurfer’s mri_vol2surf (Dale et al., 1999; Fischl et al., 1999) (http://surfer.nmr.mgh.harvard.edu/). We then resampled the Glasser et al. (2016) parcellation to fsaverage, and tallied how many studies had positive indicator values intersecting with each cortical parcel (the details of the resampling procedure are described in https://wiki.humanconnectome.org/display/PublicData/HCP+Users+FAQ#HCPUsersFAQ-9.HowdoImapdatabetweenFreeSurferandHCP, and were implemented using a custom script available at https://github.com/stobyne/Spherical-Surface-Swapper). Two studies from the subjective valuation corpus were removed because they did not contain activation peaks that overlapped with cortex, leaving a final number of 198 studies.

To test for parcels that were significantly more strongly associated with one domain than the other, we performed per-parcel chi-squared tests comparing the proportion of studies with activation in that parcel between the two domains. We permuted the study domain labels (DN or valuation) 5000 times while preserving the total number of studies in each domain, and on each iteration stored the maximum resulting chi-squared statistic across all parcels. This gave us a null distribution of 5000 maximum chi-squared values. The 95th percentile of this distribution served as an FWE-corrected significance threshold to evaluate unpermuted chi-squared values.

### 2.4. Resting-state fMRI Data

Our fMRI analyses used resting-state fMRI data from the Human Connectome Project (Van Essen et al., 2012) Q6 release (N = 100, randomly sampled from the total pool of 469 available subjects). The Washington University Institutional Review Board approved all experimental procedures, and all subjects provided written informed consent in accordance with the guidelines set by the institution. Each subject’s data was acquired over two days at Washington University in St. Louis on a Siemens CONNECTOM Skyra MRI scanner (Siemens, Erlangen, Germany). Four resting state runs (repetition time = 0.720 s, echo time = 33.1 ms, flip angle = 52°, multiband factor = 8, 72 slices, 2 mm isotropic voxels) each comprised 1200 time points (14 min 24 s) for a total of 4800 time points. Two runs were acquired on each day, with the phase encoding direction set to left-right for one run and right-left for the other. Only subjects with both left-right and right-left phase encoding for each day were included (i.e. subjects with four resting-state fMRI sessions). In addition, only datasets with low motion levels (under 1.5 mm) and less than 5% of points over 0.5 mm framewise displacement (FD; Power et al., 2014) were used. See (Van Essen et al., 2012) for more details about the data acquisition protocol.

Data initially underwent the HCP minimal preprocessing pipeline (Glasser et al., 2013), which included gradient nonlinearity correction, motion correction, EPI distortion correction, high-pass filtering (0.0005 Hz threshold), MNI152-based normalization, surface reconstruction, and mapping of functional data to a standardized cortical surface model (details can be found in Glasser et al., 2013). In addition, data underwent temporal denoising based on independent components (FMRIB’s ICA-based X-noiseifier, FIX; Griffanti et al., 2014; Salimi-Khorshidi et al., 2014). Data were further preprocessed using an in-house pipeline described previously (Tobyne et al., 2017). Steps (in order) included linear interpolation across high motion timepoints with over 0.5 mm of FD, band-pass filtering (allowed frequencies ranged from 0.009 and 0.08 Hz), and temporal denoising via mean grayordinate signal regression (Burgess et al., 2016). Interpolation of high motion time points was performed to avoid temporal smoothing of noisy signal from head motion into the filtered signal during the bandpass procedure. After filtering and denoising, the interpolated high-motion time points were censored by deletion and each run was temporally de-meaned. The processed time series had a median of 4799 time points (minimum = 4661) across participants. Each subject’s brain was comprised of 32k standard grayordinates per hemisphere (combined in a CIFTI file). We retained only the cortical surfaces, which resulted in 59,412 total surface vertices per subject.

### 2.5. Network Definition

All network analyses were performed using the igraph package (v. 1.1.2; https://igraph.org/r/; Csardi & Nepusz, 2006) in R (v. 3.4.1; https://www.r-project.org/; R Core Computing Team, 2017). To establish each subject’s network, we selected all the vertices contained within the mPFC/PCC search space (n = 4,801 per subject; mPFC = 2854, PCC = 1947) and computed the Pearson correlation of the time series for every pair of vertices. All correlation values were transformed using Fisher’s r to z. This produced a weighted network for each subject, in which the nodes were surface vertices and the edge weights were the correlations among them. Edges mostly consisted of positive correlations (mean proportion positive = 0.65, SD = 0.03). We chose not to threshold the network, as the SP algorithm is well equipped to operate on complete (i.e. fully-connected) weighted graphs (Chung, 1997). However, our results were unchanged if we retained only significant correlations (*p* < 0.05, uncorrected) in the weight matrices. Next, we took the exponential of the z-transformed correlations so that all weights became positive while maintaining their ordinal ranks. Ensuring that all edges were positive facilitated the construction of the graph Laplacian (see below), which requires all off-diagonal elements to have the same sign by design. We generated and analyzed network weight matrices at four levels: (1) for each subject’s full concatenated dataset (up to 4800 TRs); (2) on each step of a sliding window analysis (see Section 2.7 for more details); (3) for the concatenated time series for the two runs on each day (up to 2400 TRs); and (4) for each run separately (up to 1200 TRs).

### 2.6. Community Detection

Communities (i.e. clusters) were identified using the SP algorithm (Belkin & Niyogi, 2003; Chung, 1997; Fiedler, 1975; Higham et al., 2007). First, each network was represented as an *n* x *n* network weight matrix *W* as described above (where *n* equals the number of vertices in the search space, 4,801). The matrix was then transformed into its symmetric normalized Laplacian form

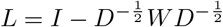

where *I* is an identity matrix of size *n*, and *D* is a diagonal matrix containing the strength of each vertex (i.e. the sum of its edge weights with all other vertices). This resulted in a matrix wherein each entry was the negative normalized value of the connection (from 0 to 1) between any two vertices relative to their combined connectivity strength, and with ones along the diagonal. The transformation ensures that every row sums to zero. We then computed the eigenvalues and eigenvectors of the symmetric normalized Laplacian matrix, and used the eigenvector associated with the second-to-lowest eigenvalue (traditionally called the ‘Fiedler vector’) to divide the network into two. The Fiedler vector consists of a set of positive and negative values and is binarized by sign to partition the network into two similar-sized communities (Fiedler, 1975). In this way, SP avoids producing communities that are too small to be physiologically meaningful (for example, small sets of vertices that are spuriously correlated due to measurement noise). Given that this data-driven method does not label the two communities or establish correspondence across participants, we defined each individual’s “DN” community as that which contained the majority of the vertices in the DN-specific PCC parcel identified in our meta-analysis (area 7m). The completeness of the graphs ensured that SP did not face the issues associated with its use in sparse networks (Fortunato & Hric, 2016).

In order to evaluate the validity of the resulting partitionings across community-detection methods, we also estimated network communities using the more traditional approach of modularity maximization (Garcia et al., 2018), based on the algorithm from Clauset et al. (2004). The method heuristically iterates through many possible combinations of vertices, and selects the partitioning that maximizes the within-community edge weights, relative to a random network containing the same number of edges and communities. Unlike SP, modularity can fractionate a network into more than two communities. Agreement between the partitions provided by the bounded (SP) and unbounded (modularity) community detection methods would suggest the results are not distorted by the restriction of SP to binary partitionings.

### 2.7. Partition Evaluation

We used the Adjusted Rand index (ARI) to evaluate the stability and topographical heterogeneity of the communities within and across individuals (Hubert & Arabie, 1985), which was calculated using the “mcclust” package in R (Fritsch, 2012). The ARI is a metric that quantifies the similarity between two alternative clusterings of the same data. The base of the ARI is computed by the formula

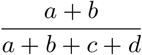

where *a* is the number of pairs of nodes that were grouped together in both partitionings, *b* is the number that were grouped separately, and *c* and *d* denote the number of pairs grouped together (separately) in one partitioning, but separately (together) in the other. Therefore, the ARI estimates the fraction of all possible node pairs that had the same status (connected or not) in both partitionings (with the denominator equal to *n*(*n -* 1)*/*2). The resulting ratio is adjusted against a baseline given by the expectation assuming independent partitionings to yield an index that ranges from 0 to 1, where 0 denotes the value expected by chance. This means that even though differences are heavily penalized, positive ARI values compare favorably against chance clustering (and the index can take negative values if the ratio given by the formula above falls below the chance level). In short, the ARI quantifies the chance-corrected agreement between any two partitions while being agnostic to the labeling scheme.

We performed a number of comparisons among partitions. First, we computed the degree of agreement between SP and modularity maximization per subject. SP and modularity maximization have been previously found to show a tendency toward underfitting and overfitting, respectively, in their community detection performance in a diverse set of network types (Ghasemian et al., 2019), so alignment between the two algorithms would increase our confidence in the validity of the resulting partitionings. Next, we compared the subject-level SP partitionings across individuals, and calculated the mean pairwise ARI for the group. We then performed the same evaluation for PCC and mPFC separately, and examined whether there were differences in overall agreement within these regions by performing a paired permutation analysis. For each individual and region we took the mean ARI with all 99 other individuals, then took the difference between regions to get an ARI difference per subject. On each of 5000 permutations each subject’s ARI difference was independently sign-flipped and the group mean difference was added to a null distribution. The unpermuted group mean difference was then evaluated against this permuted distribution.

To identify vertices whose community assignment was more stable or more variable, we performed a sliding window analysis (20 min windows, 1 min increments, median number of windows per subject = 37, range = 35 - 37), comparing each window’s resulting partitioning against the partitioning derived from the subject’s whole data set. A 20-min window has previously been found to yield relatively stable and unbiased estimates of individual-level brain network characteristics (Gordon et al., 2017). We assessed whether the magnitude of the Fiedler vector value for a given vertex (for the full subject-level data set) was associated with the stability of that vertex’s sub-network assignment across time windows. To do this, we fit a mixed effects logistic regression model, in which the dependent variable was the proportion of times each vertex participated in the DN community across windows, and the explanatory variables included a random effect of subject and a fixed effect of the Fiedler vector value for that vertex (derived from their full time series). Based on this significant relationship, we identified a threshold Fiedler vector value for each subject, such that empirical above-threshold vertices were persistently associated with either DN or non-DN more than 99% of the time.

We then estimated the level of agreement between network partitions estimated using data across individual scan days (with 2 days per participant). If the functional organization estimated by SP is indeed individual-specific, we should see higher agreement within individual (test/re-test across days) than across individuals. We tested this idea by computing the ratio of the mean ARI within and between individuals. Ratios close to one would denote similar within-participant and across-participant alignment, whereas ratios considerably higher than one would suggest that partitions were more similar within-participant than across participants. We then extended this idea by computing the agreement across individual runs (4 per subject). Similar to the day-based analysis, we assessed whether run-level data showed higher agreement within-subject than between subjects.

### 2.8. Seed-based Resting-state Functional Connectivity versus Community Detection

We evaluated the performance of the SP algorithm in comparison to a simpler partitioning approach based on seed-based functional connectivity. Independently for each day (2 per individual), we estimated each subject’s DN partition in mPFC based on its vertex-wise functional correlations (Pearson) with the spatially averaged activity across all vertices in the PCC search space. We used the whole PCC region because it is traditionally thought to be a prominent node of the DN (Buckner et al., 2008), and is a common area for researchers to place seeds for vertex-and volume-based connectivity analyses (e.g. Fox et al., 2005). We compared these seed-based maps with the unthresholded Fiedler vectors produced by SP, with the sign of the Fiedler vector oriented so the DN community was marked by positive values in every subject. We calculated three sets of across-day similarity values for each individual: 1) between the two seed-based maps; 2) between the two SP-based maps; and 3) between seed-and SP-based maps. Because the values in the maps were continuous-valued (and not categorical labels, which would be amenable to ARI), we quantified the similarity between maps in terms of the spatial Spearman correlation across vertices. These spatial correlations were meant to determine the test/re-test reliability of each approach, as well as the overall level of agreement between them. For 8 subjects, the communities produced with one of the days’ data sets had split coverage of area 7m, and our community labeling scheme for the Fiedler vector produced a sign mismatch across days. ARI is robust to such labeling issues, but the inconsistency produced strong negative correlations of the Fiedler vector across days for these individuals. Visual inspection showed that the community layout was well aligned across days, and so we matched the labeling of their partitionings based on the day that sufficiently covered area 7m.

The two methods were expected to produce somewhat similar results, but the one displaying greater within-subject agreement across days should be preferred (for a discussion on the stability of functional networks see Kong et al. (2018) and Gratton et al. (2018)). We therefore compared the within-subject spatial correlation coefficients produced by each method through a paired permutation analysis. For each of the 100 individuals, we computed the difference in inter-day correlations between methods, randomized the sign of these values 5000 times, and computed the mean of these differences on each iteration. The empirical difference in means was then evaluated against this permuted distribution.

### 2.9. Associations with sulcal morphology

Next we asked whether the location of the DN and non-DN communities was systematically related to sulcal morphology. Based on a previous report of individual alignment of DN within sulci in ventral mPFC (vmPFC; Lopez-Persem et al., 2019), we subdivided our search space into three regions: vmPFC, which matched the ROI used by Lopez-Persem and colleagues (2019; areas 25, s32, a24, 10v, 10r, p32, and OFC); superior mPFC (sup-mPFC), encompassing the remaining dorsal areas in our mPFC space; and all of the PCC search space. We used each subject’s curvature maps from the HCP (transformed to fs_LR 32k space), in which cortical depth is quantified by negative numbers for sulci and positive numbers for gyri. For each individual, we computed the Spearman correlation between curvature and the unthresholded Fiedler vector values in each region and hemisphere separately (6 correlations per individual). Since the DN was indicated by positive FV values, negative correlations meant that the DN was more likely to be contained in sulci, with non-DN located in gyri. Finally, we collected all individual correlations for each combination of hemisphere and region, and performed a one-sample t-test on each set to determine whether correlations were significantly different from 0 in our group (6 tests total).

## 3. Results

### 3.1. Meta-analysis

We performed a coordinate-based meta-analysis to identify cortical surface parcels within mPFC and PCC that were preferentially associated with the DN or with subjective valuation. Volumetric coordinates from 80 studies with task deactivation contrasts and 198 studies with valuation contrasts were projected onto a cortical surface, and mapped to discrete parcels from a multimodal cortical parcellation (Glasser et al., 2016) to produce a list of brain areas reported per study. The 40 parcels considered were limited to the medial portion of the default and limbic networks defined by the Yeo et al. (2011) 17-network parcellation. Domain-specificity was tested by first permuting the domain labels across studies (DN or valuation) to create a null distribution for the maximum chi-squared statistic in the search space (see Methods for details). The null distribution was used to identify regions that were reported significantly more often in one literature or the other.

Figure 1 shows the proportion of times each parcel was reported for each domain, as well as the significant differences between domains. The 95th percentile of the permuted chi-squared distribution was 8.87. Based on this threshold, area 7m in PCC/precuneus was the only parcel to show a preferential association with the DN bilaterally (Left: observed *χ*^2^ = 10.07, *p* = 0.029; Right: observed *χ*^2^ = 18.89, *p* < 0.001). The adjacent area v23 exhibited a similar effect, albeit only unilaterally (Right: observed *χ*^2^ = 11.51, *p* = 0.011; Left: observed *χ*^2^ = 8.25, *p* = 0.067). There appeared to be a bilateral preference toward valuation effects in mPFC area 25 (Left: observed *χ*^2^ = 12.91, *p* = 0.005; Right: observed *χ*^2^ = 12.83, *p* = 0.005); however, closer inspection suggested this effect was driven by subcortical foci centered in adjacent ventral striatum. No other parcels were preferentially implicated in valuation relative to DN. We therefore selected area 7m as an interpretable, bilateral reference point for labeling DN and non-DN communities in the analyses that follow. We note that the area labeled 7m in the parcellation used here (Glasser et al., 2016) is different from (and located inferiorly on the medial surface to) the non-DN area 7m discussed in previous work (Andrews-Hanna et al., 2010).

**Figure 1:**
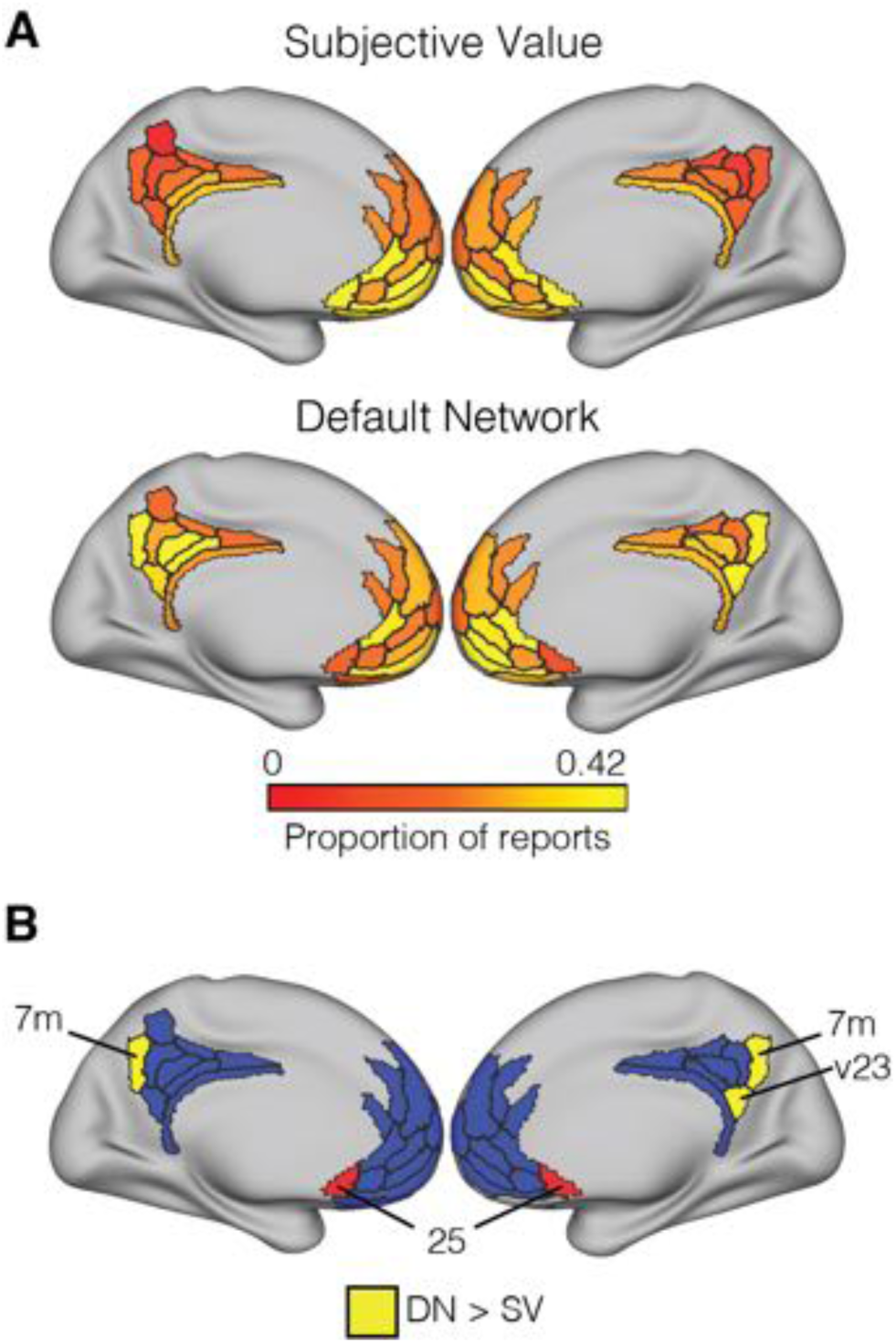
Meta-analysis results. A: Proportion of times each ROI was reported in the valuation and DN literatures. B: Regions identified in permutation-based chi-squared tests contrasting the two literatures. Area 25 (in red) initially appeared to be associated with valuation, but was not interpreted because the effect was found to reflect carryover from subcortical foci centered in ventral striatum (see text for details). Areas in blue represent the remainder of the search space.

### 3.2. Individual-level DN and non-DN communities

Within the mPFC/PCC search space, we estimated the topography of the DN for each individual. Using each individual’s full time series (approximately 4800 total TRs from four 14-min scanning runs acquired over two days), we calculated the full vertex-to-vertex correlation matrix for the 4801 surface vertices in the search space. We represented each individual’s correlation matrix in the form of a network, with cortical surface vertices as nodes and transformed correlation values as edge weights. We then applied the SP community detection algorithm to partition the network into two cohesive functional communities.

Figure 2 shows a representative partitioning of the search space for a single participant (100307; additional examples are presented in the first two columns of Supplemental Figure 1). The SP algorithm subdivides a network according to the positive versus negative values in the Fiedler vector (the eigenvector related to the second-to-lowest eigenvalue of the network’s normalized Laplacian matrix, see Methods). Since this is a data-driven approach, there is no a priori labeling for the two communities. We assigned the DN label to the community that contained the majority of the DN-specific PCC parcel from the meta-analysis (7m). We oriented each individual’s Fiedler vector so positive values corresponded to the DN community (Nenning et al., 2017), and were assigned a value of 1 in the binarized partitionings (with 0 denoting non-DN). In qualitative terms, the resulting patterns contained substantial DN coverage in posterior PCC (as dictated by our labeling strategy), with non-DN vertices in anterior PCC. The mPFC region tended to include DN vertices in its ventral-anterior and dorsal-anterior areas, with a persistent non-DN pocket between them. This non-DN section extended posteriorly into pregenual cingulate cortex (area a24). We note that the addition of restrosplenial cortex (an area commonly regarded as part of canonical DN) to the search space did not change these results; as expected, that area tended to be largely assigned to the DN community (Supplemental Figure 1).

**Figure 2:**
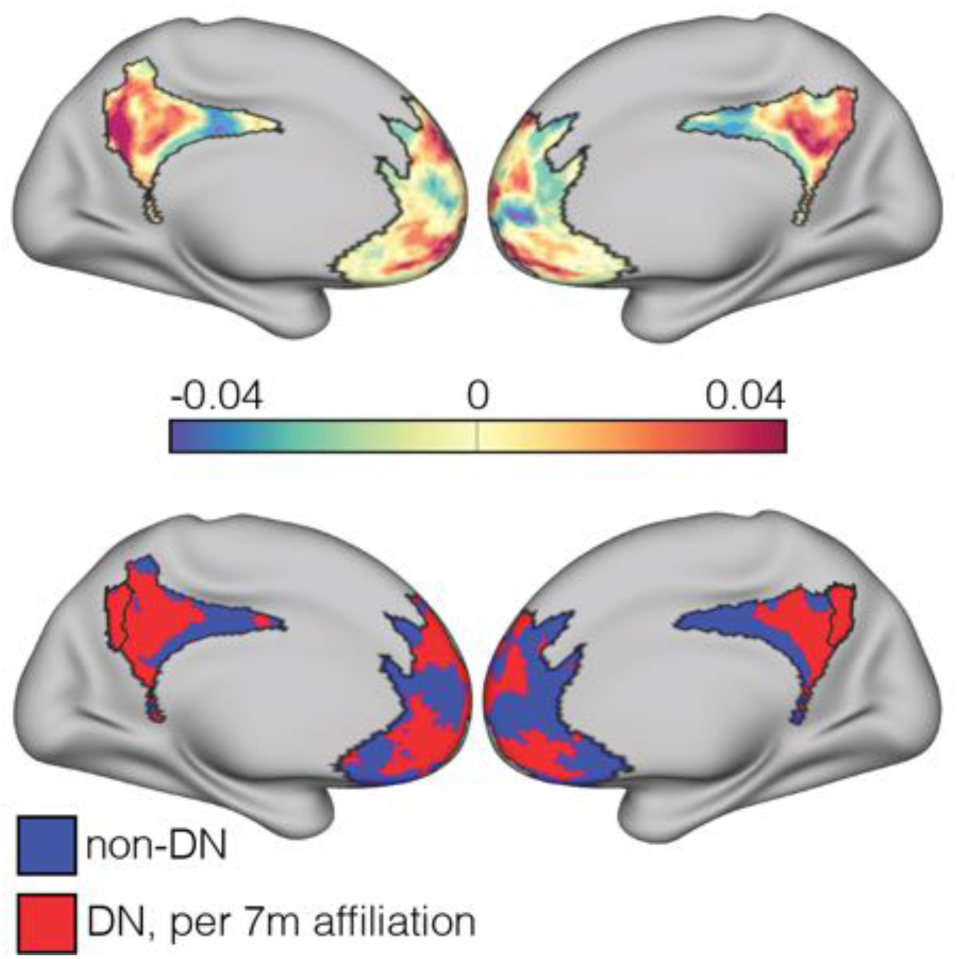
Brain partition for an example subject (100307). Fiedler vector values (top) are mapped onto the brain surface, dividing it into positive and negative communities. The bottom brain shows the binarized Fiedler vector, with red areas denoting the DN community (as indicated by coverage of area 7m, bordered).

Before evaluating the degree of generalizibility of this topographic pattern across individuals, we examined the validity of the partitionings by comparing them to results from an alternative community detection algorithm, modularity maximization (Clauset et al., 2004). Modularity seeks to find the set of communities that maximizes within-community connection weights relative to a null model. Since modularity is not constrained to a predetermined number of communities, it was capable of finding more than two in our data set. We quantified the cross-method agreement in terms of the Adjusted Rand Index (ARI; see Methods), which measures the proportion of node pairs in a network that were either clustered together or separately in both partitionings, while being agnostic to labeling schemes and controlling for chance clustering. The ARI normally takes values ranging from 0 to 1, with 0 indicating chance agreement (but can take negative values if the similarity falls below chance). Supplemental Figure 2 contains examples of ARI values in real and simulated contexts.

The two clustering methods had high agreement (mean ARI = 0.87, SD = 0.13). Modularity showed a tendency to produce additional communities (median = 3, range = 2, 5). However, the additional communities encompassed a small number of vertices (median = 16.5, IQR = 6 −41.5) compared to the principal two (median = 4783.5, IQR = 4759.5 - 4795), suggesting that a binary partitioning provided a reasonable approximation of the network’s true community structure.

Next, we examined the similarity of SP-based partitionings across individuals by computing the ARI between every pair of subjects, and found modestly above-chance agreement overall (mean = 0.13, SD = 0.05). Qualitative inspection of the community organization showed good alignment for PCC, whereas the pattern in mPFC was consistent but shifted topographically across subjects. To quantify this heterogeneity in mPFC, we calculated the between-subject ARI for each region separately (Figure 3). The functional topography of PCC was better aligned across individuals (mean = 0.19, SD = 0.09) than mPFC (mean = 0.1, SD = 0.05; paired permutation, *p* < 0.001; Cohen’s D = 1.26), although the mean ARI in mPFC still exceeded the chance value of zero (Wilcoxon signed rank test, *p* < 0.001; Cohen’s D = 2.03).

**Figure 3:**
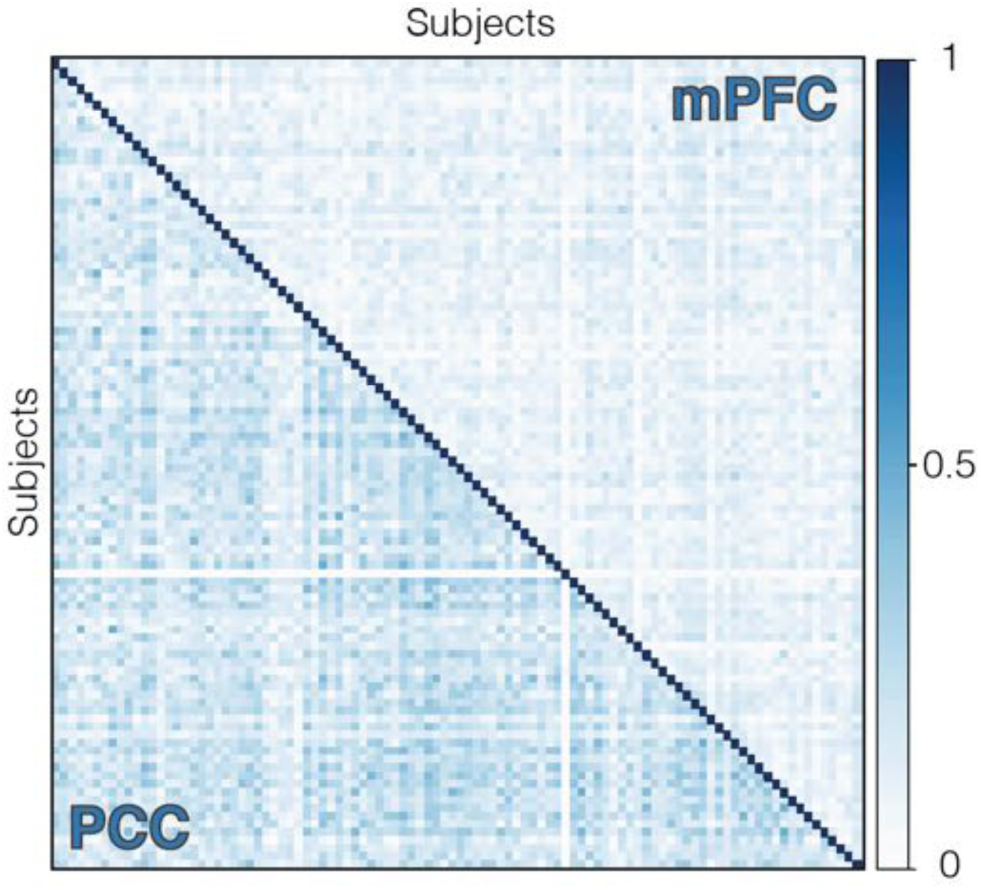
Similarity matrix showing ARI values among all subjects for PCC (lower triangle) and mPFC (upper triangle) separately. Functional topographic patterns were better aligned across individuals in PCC than mPFC.

### 3.3. Pattern variability over time

We next sought to estimate whether individual vertices had a stable or unstable community affiliation over time. We did so by performing a sliding window analysis on each subject’s full time series (20 min windows shifting by 1 min). We compared the partitioning derived from each window with the partitioning computed using the entire time series (Figure 4). Our focus here was not on the overall level of agreement (which is expected to be high given the use of overlapping data), but on differences in stability across nodes. The sliding window analysis provided a means to identify nodes that were highly variable, and allowed us to determine whether these variable nodes followed a specific spatial structure.

**Figure 4:**
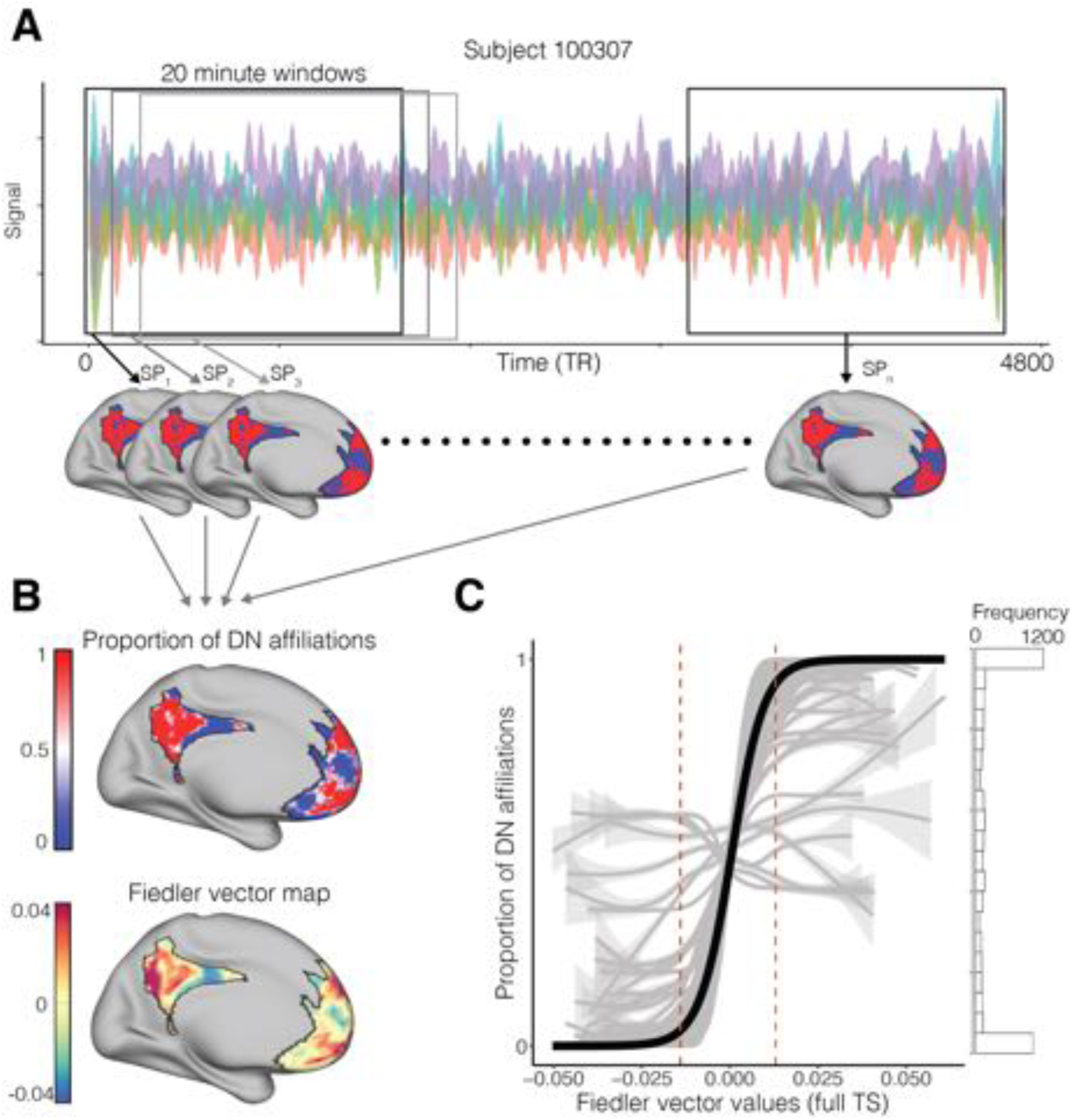
A: For each individual, we produced partitions for each 20 minute sliding window (84 TRs). B: Proportion of times each vertex was affiliated with the DN community across windows in one example subject (upper), and the continuous Fiedler vector map for the same subject using their full time series (lower). C: Relationship between the magnitude of Fiedler vector values and the proportion of DN affiliations. Grey lines display data for each subject, and the black line shows the fit from a mixed-effects logistic regression. Dashed red lines indicate the mean FV value at which maps were thresholded. The histogram displays the mean frequency distribution of y-axis values.

The mean ARI along each subject’s time series was significantly higher for PCC (mean = 0.59; SD = 0.14) than mPFC (mean = 0.5; SD = 0.13; paired permutation, *p* < 0.001; Cohen’s D = 0.65). A subset of nodes showed exceptionally high stability, in that they were assigned to the same community in every time window. The percentage of stable nodes ranged from 0 to 73% across individuals (median = 49.5%, IQR = 29% - 60.25%).

We next tested whether the continuous-valued Fiedler vector (before binarization into discrete communities) carried information about the stability of individual nodes. There is precedent in the literature for the idea that the magnitude (and not just the sign) of the Fiedler vector values conveys important information about the role of each node in the network (Gkantsidis et al., 2003; Tian & Zalesky, 2018). Therefore, we tested whether the magnitude of the eigenvector values was associated with the stability of nodes over time. Specifically, we estimated the proportion of DN affiliations per node as a function of Fiedler vector values, using a logistic mixed effects model (Figure 4). The model identified a positive significant relationship between these features (*β* = 217.02, SE = 0.67, *p* < 0.001), signifying that vertices with higher absolute Fiedler vector values were more persistent in their relationship with their corresponding community over time.

These analyses suggest that there is potential value in thresholding the Fiedler vector as a means to identify reliable DN and non-DN vertices on an individual subject basis. We therefore thresholded each subject’s Fiedler vector to produce these refined maps. For each individual, we estimated the threshold by selecting the empirical smallest absolute Fiedler vector value that yielded an average stability across suprathreshold nodes of 99%, for positive (mean = 0.0132, SD = 0.006) and negative (mean = −0.0139, SD = 0.0069) values separately. Individuals without such stable nodes (n = 19) were not thresholded, and were included in the subsequent analyses in unthresholded form. The median proportion of retained vertices per individual was 0.49 (IQR = 0.29 - 0.65). Sub-threshold vertices were set to zero in Fiedler vector maps and 0.5 in the binarized maps (so that they would not bias the calculation of averages). Figure 5A shows the thresholded partitioning for the same individual shown in Figure 2. The maps used in all subsequent analyses were thresholded by this individualized criterion.

**Figure 5:**
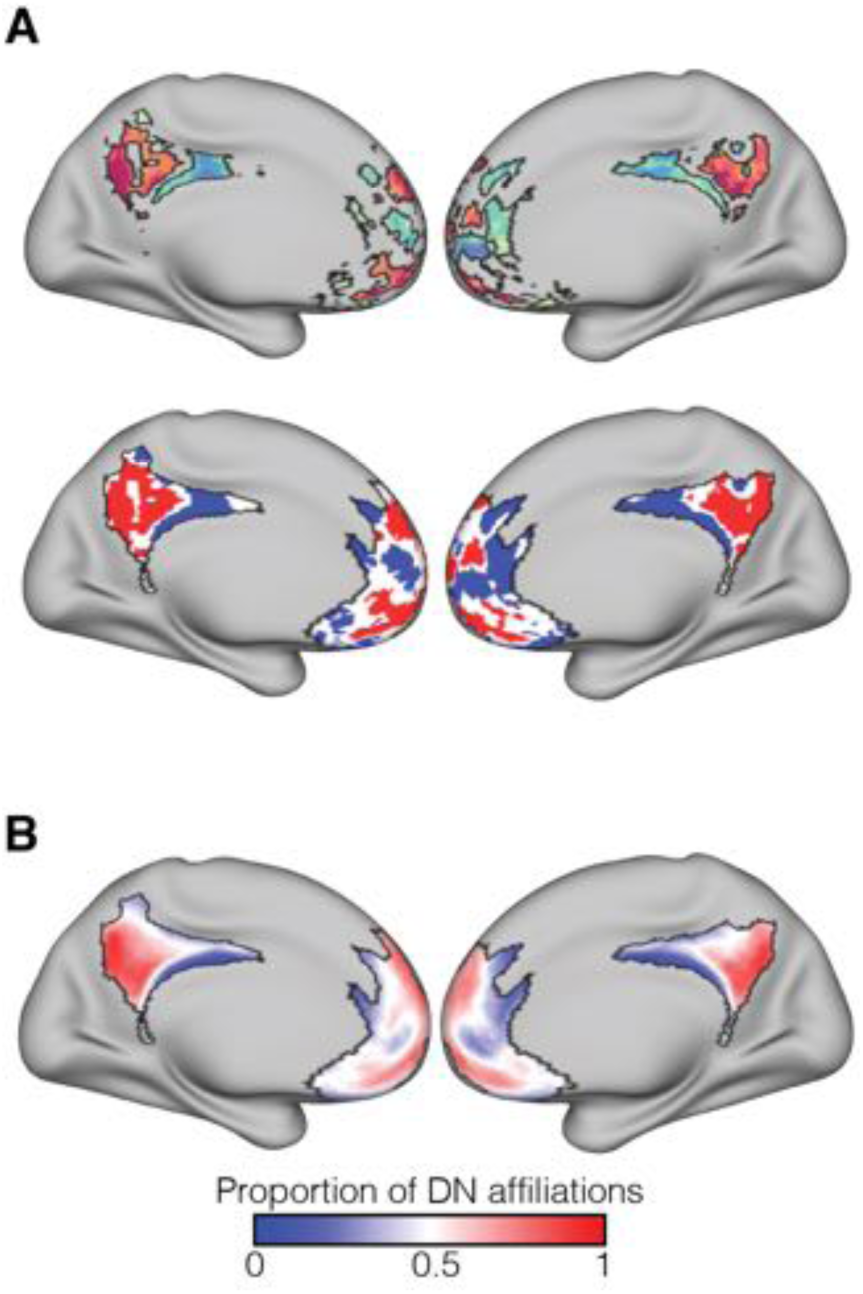
A: Thresholded Fiedler vector map for subject 100307 (top), and its binarized form (bottom). Subthreshold values effectively formed a third community of high-variability vertices. B: Mean of the binarized maps across all participants, indicating the proportion of DN affiliations per vertex in our sample. Colors represent PCC-based labels (‘DN’ versus ‘non-DN’), which were applied in a subsequent step following the data-driven community-detection analysis and which were necessarily well-aligned in PCC. This aggregate map shows the common organizational principle of the DN and non-DN communities, while also showing the high level of variability in mPFC.

With these thresholded partitions, we recomputed the overall similarity across participants. Compared to before, there was lower topographic agreement across individuals (mean ARI = 0.07, SD = 0.04). The same was true for both PCC (mean = 0.1, SD = 0.07) and mPFC (mean = 0.05, SD = 0.03) separately, although the significance of the differences between areas was preserved (paired permutation, *p* < 0.001; Cohen’s D = 1.11). Figure 5B shows the average of the thresholded partitions across all participants, denoting the proportion of times a vertex was affiliated with the DN community. This summary illustrates the common organizational layout of both communities, but also highlights the considerable variability across individuals. To test the possibility that the higher inter-subject variability in mPFC was driven merely by lower signal quality in the retained vertices, we quantified the temporal signal to noise ratio (tSNR) for each region, both before and after thresholding. We calculated tSNR using time series that were not demeaned, but were otherwise equivalent to the data originally used. A map of the mean tSNR across individuals can be found in Supplemental Figure 3. In terms of tSNR variability across vertices within each region, mPFC had overall greater spatial standard deviation both before and after thresholding (mPFC: pre-threshold mean spatial SD = 33.96, post-threshold mean spatial SD = 30.15; PCC: pre-threshold mean spatial SD = 15.28, post-threshold mean spatial SD = 14.59). However, mean tSNR after thresholding was significantly higher for mPFC than PCC (mPFC: mean = 77.34, SD = 13.77; PCC: mean = 64.99, SD = 10.19; permutation p-value < 0.001, Cohen’s D = 1.02). This reflected a significant increase in mean tSNR in mPFC as a result of the thresholding step (pre-threshold mean = 66.5, SD = 7.87; paired permutation p-value < 0.001, Cohen’s D = 0.97), whereas the mean signal quality in PCC increased only slightly (pre-threshold mean = 64.56, SD = 10.02; paired permutation p-value = 0.0384, Cohen’s D = 0.04). In short, mPFC had higher overall tSNR, albeit with greater variability across nodes. Applying the thresholding step focused the analysis on vertices with high signal quality.

### 3.4. Test/re-test reliability across days

The relatively high inter-individual variability seen in the aggregate map could reflect at least three factors: (1) measurement noise, (2) dynamic variation in mPFC network organization, and (3) stable patterns of functional organization that differ across individuals. To arbitrate among these possibilities, we examined the test/re-test reliability of mPFC/PCC community structure across separate days of testing. Insofar as the observed variability reflects individual-specific brain organization, across-day ARI values should be consistently higher within-individual than between individuals (an example comparison for two individuals is provided in Supplemental Figure 2). Figure 6 shows pairwise comparisons among ten example subjects for PCC and mPFC separately (left).

**Figure 6:**
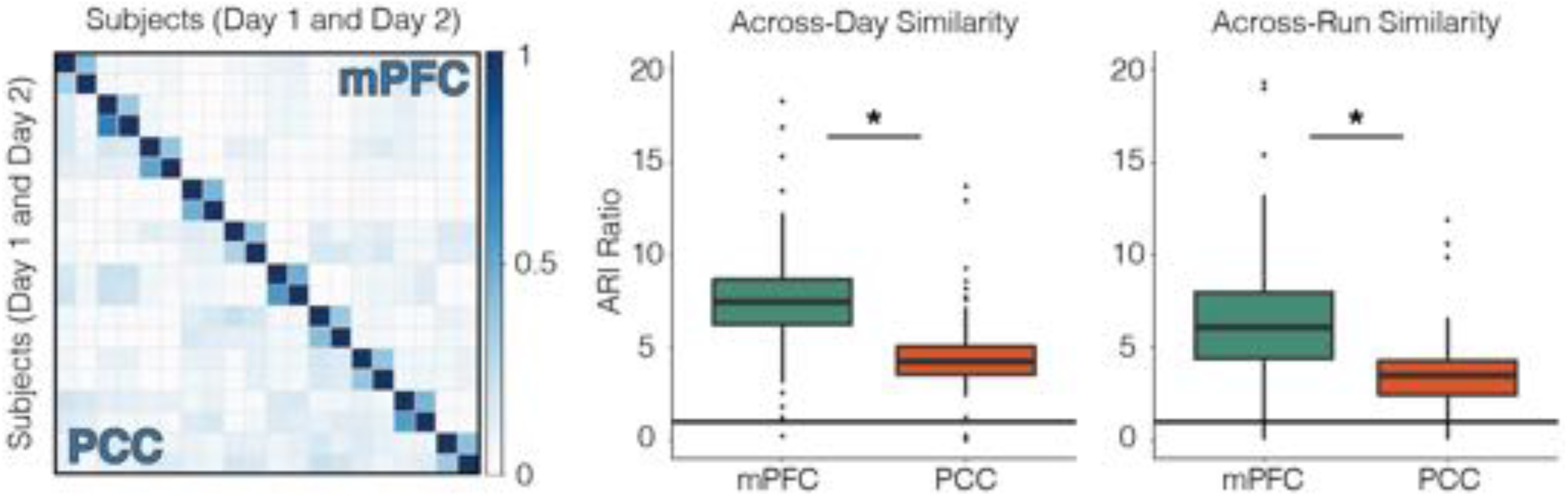
Left: Similarity matrix for 10 example participants (2 scanning days each), showing pattern agreement across days and subjects for PCC and mPFC separately. Color scale represents the ARI, which quantifies topographic similarity irrespective of how the communities are labeled. The block-diagonal structure is indicative of test-retest reliability across days within an individual. Middle: ratio of within-subject ARI to between-subject mean ARI for all individuals across days suggests idiosyncratic community arrangement for both PCC and mPFC (ratios > 1, solid line), with greater subject-specificity in mPFC. Right: within-to-between subject mean ARI ratios for run-specific partitionings again show greater subject-specific organization for mPFC.

Once again we found low alignment across individuals for PCC (mean = 0.08, SD = 0.06) and mPFC (mean = 0.03, SD = 0.03), but both areas showed comparatively high levels of within-individual agreement (PCC: mean = 0.36, SD = 0.14; mPFC: mean = 0.26, SD = 0.1). We calculated an index of relative specificity by computing the ratio of each individual’s across-day (within-participant) ARI to the mean of all between-participant ARI values involving that individual. The index is expected to take on a value near 1 if partitionings are well aligned across individuals and/or are subject to a common level of measurement noise. It is expected to exceed 1 insofar as functional network organization is reliable and individual-specific. This index is intended to factor out the potential contributions of measurement noise or dynamic instability, which would introduce variability both across individuals and across days.

Figure 6 shows ARI ratios for PCC and mPFC. A signed-rank test showed evidence for specificity (i.e. ratios > 1) in both mPFC (median = 7.45, IQR = 6.08 - 8.65, V = 5037, *p* < 0.001) and PCC (median = 4.25, IQR = 3.53 - 5.29, V = 5030, *p* < 0.001). Moreover, the ratios for mPFC were significantly greater than those for PCC when compared in a paired permutation test (*p* < 0.001; Cohen’s D = 0.5). This pattern was mostly unchanged when computed using modularity maximization to detect the communities, showing that the results persisted even without a forced binarization (mPFC: median = 4.46, IQR = 3.55 - 5.33; PCC: median = 2.85, IQR = 1.94 - 3.38; difference: *p* < 0.001; Cohen’s D = 0.52). These test/retest results suggest that the topographic variability seen in mPFC arose at least in part from stable and subject-specific organizational patterns (examples of these partitionings can be found in Supplemental Figure 4). We stress that our similarity metric, the ARI, measured the similarity of partitionings in a label-agnostic manner. The greater inter-individual consistency in PCC was therefore not merely an artifact of having used a PCC subregion as the basis for label assignment.

### 3.5. Test/re-test reliability across runs

We extended the analysis of per-day data by examining whether the organization of the DN could be extracted using per-run data only. The duration of each run (approximately 14 min) falls well below a previously suggested stability threshold for fMRI-based modularity estimations (Gordon et al., 2017). Nonetheless, high ARI ratios could indicate that the SP algorithm can still obtain information about individual-specific patterns of DN organization from a single run of data.

Run-specific SP results captured unique organizational patterns to some degree, even though the overall levels of agreement decreased (PCC between subjects: mean = 0.04, SD = 0.05; mPFC between subjects: mean = 0.01, SD = 0.02; PCC within subjects: mean = 0.17, SD = 0.14; mPFC within subjects: mean = 0.09, SD = 0.08). Supplemental Figure 4 shows that even though the community estimates were indeed less reliable within-individuals than those captured using per-day data (and sometimes even failed to produce meaningful partitionings), the layout of DN and non-DN was still observable in many cases, and was comparable to the organization seen using larger amounts of data. We again computed each subject’s ARI ratio in order to quantify the specificity of the partitions, this time using the mean of 6 across-run (within-participant) ARI values in the numerator of the ratio (Figure 6, right).

As before, a signed rank test showed that both regions had ARI ratios significantly greater than 1 (mPFC: median = 6, IQR = 4.14 - 7.99, V = 4953, *p* < 0.001; PCC: median = 3.51, IQR = 2.4 - 4.26, V = 4971, *p* < 0.001), and ratios for mPFC were higher than those of PCC (permutation *p* = < 0.001; Cohen’s D = 0.94). This result further confirms that the intrinsic functional organization of mPFC is uniquely arranged per individual, and provides evidence that information about such patterns can be extracted from relatively small amounts of data.

### 3.6. Correlation vs community detection in mPFC

We next explored the possible advantage of community detection relative to a more conventional seed-based functional connectivity analysis for estimating the individual-specific functional topography of mPFC. We examined whether maps generated with SP were more similar per participant across days than those computed from seed-based correlations. We generated a seed time-series by averaging all vertices in the PCC region of our search space, and calculated its correlation with the activity of each vertex in mPFC. The use of the whole PCC region (instead of just 7m) was meant to represent a typical approach to seed-based connectivity that relies on the group-average location of canonical DN regions. We compared the map of correlation values in mPFC to the map of unthresholded Fiedler vector values using Spearman correlations across vertices. Pairwise spatial correlations were calculated among maps computed for each day and method from all individuals. Figure 7A shows that these pairwise comparisons resembled those from the across-day comparisons above, and suggested good alignment between methods, but particularly high agreement within subject and method.

**Figure 7:**
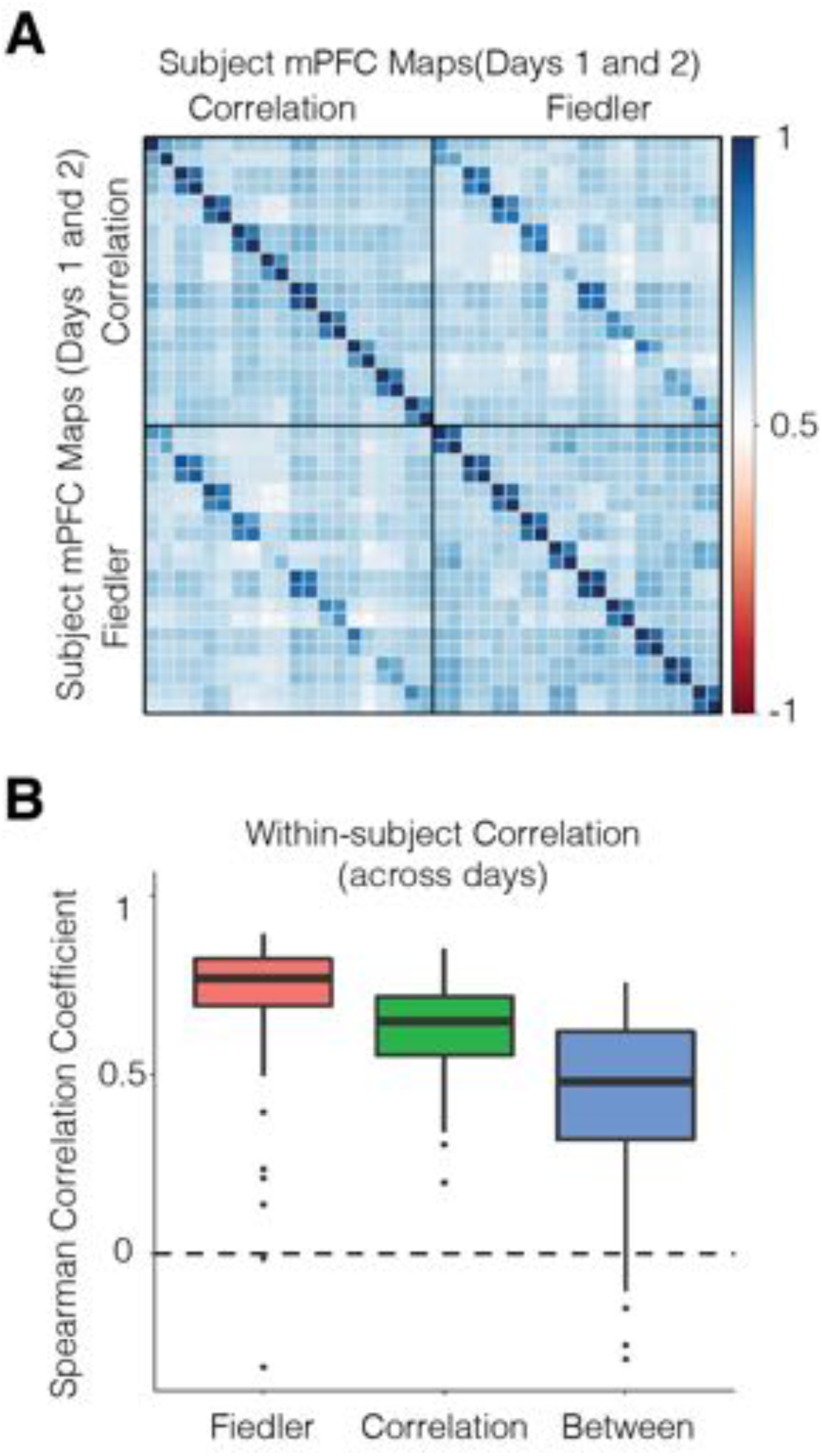
A: Correlation matrix comparing the across-day spatial stability of mPFC maps derived from seed-based functional connectivity (using a PCC seed) and the Fiedler vector for 10 example subjects. The top-left quadrant represents seed-based FC maps, and the bottom-right the Fiedler vector, with two single-day-based maps per individual. The upper-right and lower-left quadrants show across-method agreement. B: Day 1 vs Day 2 within-subject correlation coefficients for each method, as well as between methods. Community detection through spectral partitioning provided more stable estimates, even though both methods showed good levels of agreement.

Figure 7B shows the test/re-test reliability across days for patterns derived using community detection, seed-based correlation, and across methods (e.g. Day 1 community detection versus Day 2 seed-based correlation). While both approaches were reliable, community detection displayed a significantly higher median correlation coefficient across days than seed-based correlation (Community: median = 0.77, SD = 0.19; Seed-based: median = 0.63, SD = 0.12; paired permutation *p* < 0.001; Cohen’s D = 0.54). Agreement across methods was fair (median = 0.48, SD = 0.23), signifying that the two approaches identified similar topographic features but also had systematic differences. These findings suggest that graph-theoretic community detection algorithms are advantageous for detecting stable functional topologies, in addition to their other advantages of being data-driven, unbiased and observer agnostic.

### 3.7. Relationship between functional organization and sulcal morphology

Next, we asked if the idiosyncratic organization of the DN corresponded to patterns of sulcal morphology. Several previous studies have provided evidence that sulcal and gyral organization informs the location of functional effects (Amiez & Petrides, 2014; Amiez et al., 2013; Zlatkina et al., 2016). Recent work has suggested that DN regions in individuals lie mostly within sulci in vmPFC (Lopez-Persem et al., 2019). We sought to reproduce this relationship using the SP communities, and interrogated whether it persisted in PCC and more superior mPFC regions.

Figure 8A shows a qualitative comparison between the thresholded DN and non-DN communities and curvature maps for two individuals. In agreement with findings from Lopez-Persem and colleagues (2019), the DN community appeared to overlap with the superior rostral sulcus in vmPFC in these individuals, whereas the non-DN community included both gyri and sulci. A similar trend was observable in left PCC, where the DN community traced sulcal layouts and non-DN was more likely to appear in gyri.

**Figure 8:**
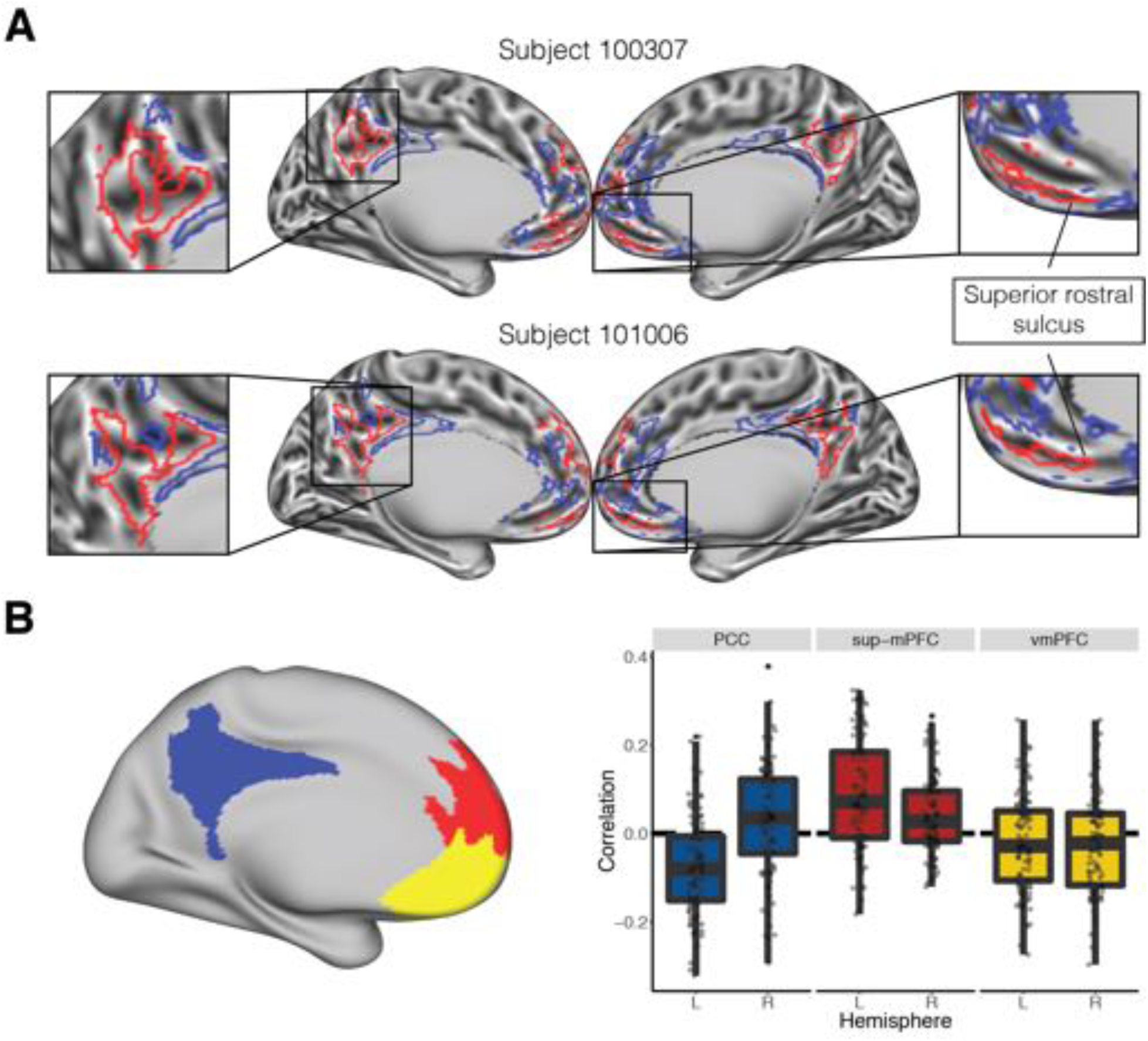
A: Correspondence between SP communities and sulcal morphology in two example individuals. Zoomed-in boxes show areas in PCC and ventral mPFC where the DN community appears to follow sulci. The superior rostral sulcus aligned with the DN community, in agreement with findings by Lopez-Persem et al. (2019). B: Correlations between cortical curvature and the unthresholded Fiedler values. We divided the search space into three areas (left): vmPFC (yellow), superior mPFC (sup-mPFC, red), and PCC (blue). Boxplots on the right show the distribution of correlation values across individuals for each combination of region and hemisphere, indicating that the DN (defined by positive FV values) was associated with sulci in vmPFC, but with gyri within sup-dmPFC. PCC showed evidence of an opposite association across hemispheres.

We quantified these observations by dividing the search space into three regions: a ventral mPFC area corresponding to the region tested by Lopez-Persem et al. (2019), a superior mPFC area (sup-mPFC) that contained the remaining mPFC regions, and PCC (Figure 8B, left). For each individual, we correlated the unthresholded Fiedler vector with cortical curvature for each combination of region and hemisphere. Negative correlations in this context imply that the DN was found in sulci and non-DN in gyri. Results are shown in Figure 8B. Correlations tended to be slightly negative in vmPFC, both on the left (mean correlation = −0.02, SE = 0.01) and right (mean correlation = −0.02, SE = 0.01); the distribution was only significantly different from zero in the right hemisphere, and weakly so (one sample t-test: t = −2.18, *p* = 0.031, uncorrected; Cohen’s D = −0.22). Correlations between FV and curvature were positive and significantly different from zero in sup-mPFC, both on the left (mean correlation = 0.09, SE = 0.01; one sample t-test: t = 6.95, *p* < 0.001; Cohen’s D = 0.69) and right (mean correlation = 0.04, SE = 0.01; one sample t-test: t = 4.49, *p* < 0.001; Cohen’s D = 0.45). Correlations in PCC were negative and significantly different from zero in the left hemisphere (mean correlation = −0.08, SE = 0.01; one sample t-test: t = −6.89, *p* < 0.001; Cohen’s D = −0.69), but significantly greater than zero in the right hemisphere (mean correlation = 0.03, SE = 0.01; one sample t-test: t = 2.19, *p* = 0.031; Cohen’s D = 0.22). The difference across hemispheres was significant (paired t-test: t = −7.07, *p* < 0.001; Cohen’s D = −0.88). These results provide preliminary indications that the association between function and structure is heterogenous across subregions of the canonical DN.

### 3.8. Alignment of mPFC community structure with a proposed DN sub-network organization

The thresholded partitions we identified had conceptual and topographic similarities to DN sub-networks A and B proposed by Braga and Buckner (2017). We explored the relationship between the two sets of sub-regions by reproducing the previously described seed-based connectivity approach in two of our subjects. In previous work, Braga and Buckner (2017; Braga et al., 2019) manually selected individual vertices in dorsolateral prefrontal cortex (DLPFC) that produced two spatially anticorrelated, interdigitated networks with distinctive patterns in the temporo-parietal junction (TPJ), inferior parietal lobule (IPL), parahippocampal cortex, mPFC, and PCC. We hypothesized that if the SP communities corresponded to one or both of the previously proposed sub-networks, our partitionings should match networks A and B generated by seed-based functional connectivity in these diagnostic areas. For whole-brain functional connectomes from two individuals (100307 and 101006), we evaluated seeds in each diagnostic region that reproduced networks A and B (correlation coefficients thresholded at 0.2), and confirmed their placement based on functional connectivity patterns observed in the remaining areas. In both individuals, seeds in posterior IPL and TPJ most clearly identified networks A and B, respectively. The whole-brain seed-based functional connectivity maps for the two individuals are juxtaposed with the corresponding community detection results in Figure 9. It is worth noting that a few distinguishing features are missing due to below-threshold correlation values (e.g. network B in right PCC of 100307).

**Figure 9:**
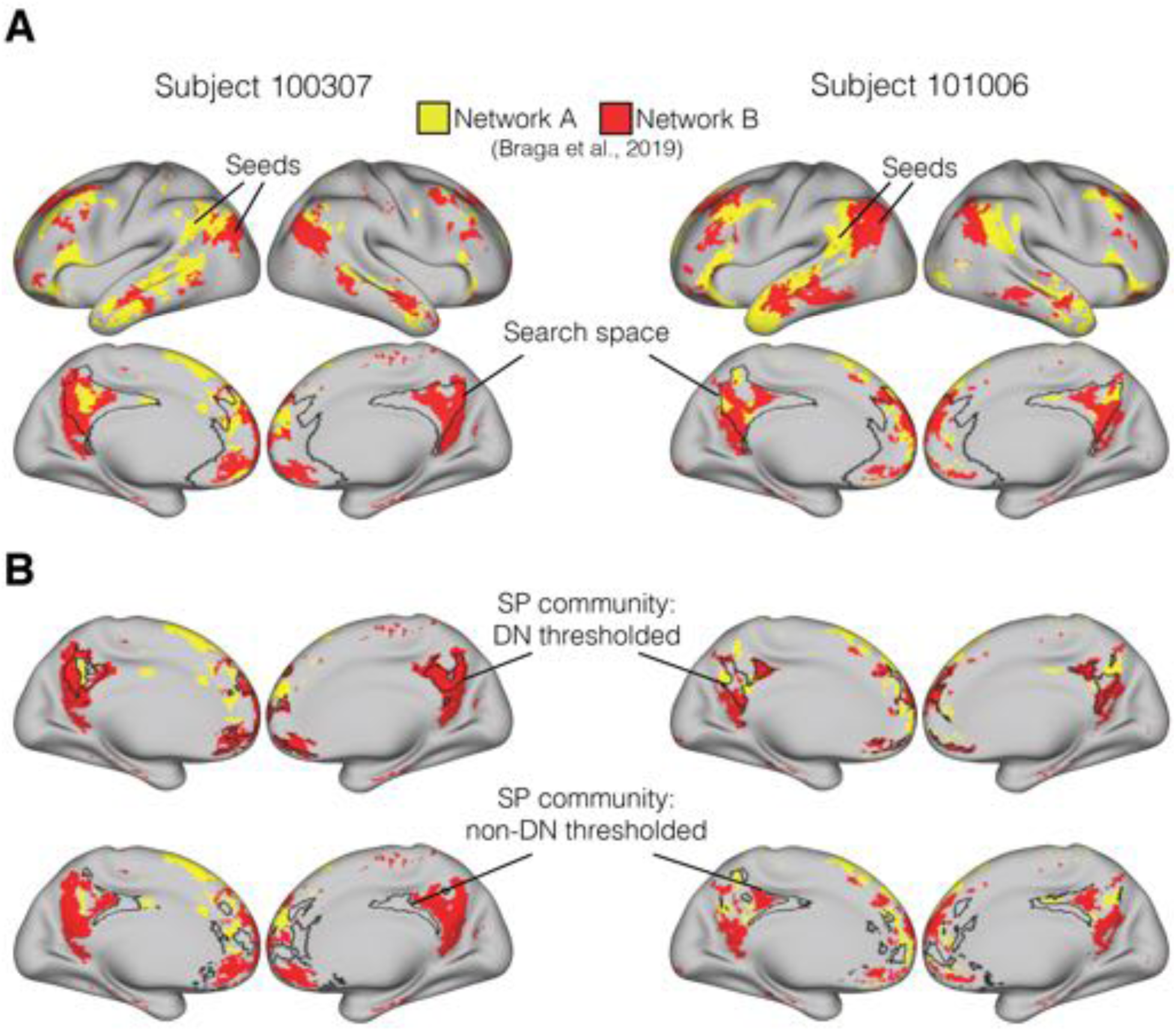
Qualitative comparison between DN sub-networks A and B (estimated based on descriptions from Braga et al. (2019)) and SP communities for two individuals. Panel A: Whole-brain networks A and B produced by selecting seeds in TPJ, with our community detection search space delineated by black borders. Correlation values are thresholded at 0.2. Panel B: thresholded communities (indicated by borders) show strong resemblance between the DN community and network A. The non-DN community covers sections of cortex not associated with either DN sub-network.

Visual inspection of these networks showed high similarity between our DN community and the previously reported sub-network A. However, the non-DN community filled areas not covered by either DN-A or DN-B. Since this three-network configuration is at odds with the two-network solution suggested in our previous analyses (i.e. comparison with modularity), we ran additional evaluations to confirm its existence. First, we reproduced the whole-brain k-means clustering analysis (12 clusters, 100 iterations) performed by Braga et al. (2019) using the full time series for two subjects (Supplemental Figure 5A). In addition to identifying DN networks A and B through this approach (in line with previous findings), we found a third cluster that aligned well with the non-DN community. To understand why modularity maximization did not identify the same three discrete clusters within the search space, we performed a silhouette analysis to determine the ideal number of clusters in our search space (Supplemental Figure 5B). For all 100 individuals, we ran k-means clusterings restricted to the mPFC/PCC search space with a specified number of clusters ranging from 2 to 5, and computed a silhouette score for each solution (higher silhouette scores indicate a better fit). Scores decreased as the number of clusters increased beyond 2 for all individuals, suggesting that a bisection was indeed the best solution. Paired permutations comparing silhouette scores across individuals indicated that the difference between two (mean = 0.042, SE = 0.001) and three (mean = 0.032, SE = 0.0009) cluster solutions was significant (*p* < 0.0001, Cohen’s D = 0.96). Visualization of a two-cluster k-means revealed a close match with partitionings estimated through modularity, and are comparable to those produced by SP prior to thresholding (Supplemental Figure 5A, bottom rows). These analyses suggest that SP isolated DN-A and the non-DN community as the dominant opposite signals within our search space, but that DN-B is observable once we take advantage of the continuous information contained in the Fiedler vector.

These results support the idea that the two approaches serve complementary purposes. Whereas Braga and colleagues (2017; 2019) identified subdivisions within the DN, the present community detection approach might be better understood as partitioning DN from non-DN cortex. Furthermore, the results show that the continuous-valued output of our approach provides a method for estimating the location of the three networks in PCC and mPFC that is less computationally demanding than full-brain clustering.

## 4. Discussion

A considerable amount of meta-analytic work has been dedicated to characterizing the brain activity patterns associated with psychological processes in medial prefrontal cortex (mPFC), revealing both dissociable and overlapping activation across domains (De La Vega et al., 2016; Hiser & Koenigs, 2018; Kragel et al., 2018). For example, topographic patterns associated with subjective valuation and with the default network (DN) have been suggested to be indistinguishable in mPFC, with overlap also partially extending to posterior cingulate cortex (PCC) (Acikalin et al., 2017; Bartra et al., 2013; Clithero & Rangel, 2014; Laird et al., 2009). This apparent overlap of task-related effects with DN regions has important implications, as it has motivated theoretical proposals about ways in which these superficially dissimilar domains might involve a shared set of core cognitive processes (Acikalin et al., 2017; Clithero & Rangel, 2014; Northoff & Hayes, 2011).

However, the interpretation of overlap in group-level data depends on the degree to which functional organization is heterogeneous across individuals. Recent studies have shown that heteromodal brain regions have considerable variability in functional connectivity across individuals (Mueller et al., 2013), individual-specific functional topography can be occluded in aggregative estimations (Braga & Buckner, 2017; Gordon et al., 2017; Michalka et al., 2015; Tobyne et al., 2018), and overlap in functional activation can vanish with increases in spatial precision (Woo et al., 2014). These findings suggest that group-level and meta-analysis-level overlap does not necessarily imply overlap in individual brains. To date, our understanding of the individual-level heterogeneity in the functional topography of mPFC has been mostly descriptive (Braga & Buckner, 2017; Braga et al., 2019; Gordon et al., 2017). A strong test of the overlap between task-related effects and DN regions would require a method to reliably and precisely capture the functional topography of mPFC in isolated individuals, as well as a quantitative estimate of the degree of topographic heterogeneity across a large group of individuals.

Here we address these challenges by using spectral partitioning (SP), a graph-theoretic community detection algorithm that efficiently separates a network into two (Fiedler, 1975; Higham et al., 2007; Toker & Sommer, 2019). For each of 100 individuals, we subdivided canonical DN regions into DN and non-DN communities. Restricting our analyses to a general mPFC/PCC search space made it appropriate to use a technique that identified a vertex-wise, binary partitioning that was sensitive to the complex topography of the brain. This contrasts with whole-brain network analyses, which need to allow for multiple sub-networks and which often use parcels that are several orders of magnitude larger than vertices as the units of analysis. Partitioning an individual’s brain network through SP has a number of advantages, including identifying communities deterministically, constraining communities to contain a similar number of vertices (i.e. preventing the allocation of most vertices to a single community), providing continuous values that relate to the strength of a node’s community affiliation, and the ability to diagnose the connectedness of a network through examination of its resulting eigenvalues (Chung, 1997; Higham et al., 2007). Comparisons with partitionings formed by modularity maximization, which heuristically determines the ideal number of communities (Garcia et al., 2018), as well as a silhouette analysis, suggested the binary partitioning was appropriate.

We found a generalizable pattern across individual partitionings, in which the DN community covered ventral and anterior/superior mPFC and posterior PCC, with the non-DN community concentrated in pregenual ACC and anterior PCC. The precise spatial positioning of this general community structure was highly heterogenous across individuals, yet stable across test/re-test evaluations within-individual. The idiosyncrasy in functional topography was particularly pronounced in mPFC, and was identified in both run-based and day-based analyses. Individual-specificity could theoretically arise from a variety of sources. For example, individual variability could be due to shifts in functional organization that are independent of structural features (Conroy et al., 2013; Nenning et al., 2017), or could relate to the pattern of functional connections with the rest of the brain (Mars et al., 2018; Passingham et al., 2002; Tobyne et al., 2018). Alternatively, the functional topography of mPFC could be governed by its underlying sulcal and gyral organization, which has been shown to vary systematically across individuals (Mackey & Petrides, 2014). Our results offer some support for this idea, echoing previous findings that DN is contained within sulci (in particular the superior rostral sulcus) in vmPFC (Lopez-Persem et al., 2019). Structure/function associations were heterogeneous in other regions; DN tended to be located in gyri in more superior mPFC regions, whereas the association differed across hemispheres in PCC. Future studies should further characterize this heterogeneous relationship. Another important goal for future work will be to assess whether the network layout in these regions can also be predicted on the basis of other aspects of brain structure, such as myeloarchitecture (Glasser et al., 2016) or structural connectivity (Osher et al., 2016; Saygin et al., 2011, 2016).

Network-partitioning methods such as SP are data-driven, and therefore provide no labeling information about the resulting communities. We circumvented this issue by independently identifying the DN community based on its coverage of area 7m, a region in PCC that was preferentially associated with the DN relative to subjective valuation in our meta-analysis. We were able to apply labels derived from this group-level approach on the basis of the topography in PCC, where functional organization was more consistent across individuals. Because each community spanned both mPFC and PCC, the labels extended to mPFC where topography was more heterogeneous.

Our results extend previous work that described individual-specific brain organization. Several recent investigations have identified topographic heterogeneity using a different data aspect ratio than we used here (a small number of individuals and a large number of scanning sessions per individual; Braga & Buckner, 2017; Braga et al., 2019; Gordon et al., 2017). Previous work has also shown that functional correlations among pre-defined cortical parcels are highly stable within an individual (Gratton et al., 2018; Kong et al., 2018). Here we were able to quantify the variability and stability of functional topography in a large sample at a fine, vertex-level spatial granularity, using moderately low amounts of data (down to a single 14 minute scan, although estimates based on more data were more reliable). The motivation to subdivide DN also stems from recent work by Kernbach et al. (2018), which identified specialized communication of parcels within DN with the rest of the brain in a large pool of individuals.

In addition to the technical advantages noted above, the SP algorithm offers analytical advantages specific to neuroscience. We found that SP outperformed a traditional seed-based correlation approach in capturing idiosyncratic functional topography. Community detection methods such as SP are stabilized by relying on all pairwise correlations among cortical vertices (rather than correlations with an individual seed). In addition, we found we could threshold the underlying Fiedler vector on the basis of the temporal stability of SP results. The magnitude of Fiedler vector values has been recently used to characterize the continuous connectivity profile of the insula with the rest of the brain, challenging the notion of discrete parcellations in that region (Tian & Zalesky, 2018). The combination of discrete classification and graded information yielded by SP provides additional flexibility and richness relative to some other clustering algorithms.

The community organization of PCC and mPFC was congruent with DN sub-networks A and B proposed by Braga & Buckner (2017; Braga et al., 2019). The topography of our thresholded DN community closely matched network A, whereas our non-DN community included cortical territory that was not part of either DN network. Subthreshold vertices from the SP communities in turn overlapped with DN-B vertices. Our findings therefore complement the initial identification of DN sub-networks by quantifying the systematic variability of their underlying topography in a larger group of people. Understanding the interaction of networks DN-A, DN-B, and non-DN is an important goal for future research. SP is related to methods that have gained traction recently for distinguishing functional cortical gradients (Huntenburg et al., 2018; Margulies et al., 2016; Tian & Zalesky, 2018). A valuable goal for future work would be to assess whether DN-A and DN-B form part of a gradual information processing sequence, or if their functions can be discretized. Regardless, this set of results collectively suggests that canonical DN regions can be topographically partitioned into DN and non-DN communities, and that the DN community can in turn be further divided into sub-networks A and B.

### 4.1. Conclusion

Our findings show that the functional topography of mPFC is variable across a large pool of individuals, and that the SP algorithm is a useful tool for identifying individualized topography in a data-driven way. The ability to capture an individual’s functional topography without the need for group priors is clinically relevant, as it could help target the assessment of mPFC activity changes in disorders such as depression and schizophrenia (Hiser & Koenigs, 2018). It will be beneficial for future task-based fMRI experiments to be able to characterize where task-evoked activity is situated relative to an individual’s overall mPFC organization. Our work is relevant to interpreting the overlap of DN regions with task-related brain activity in numerous cognitive domains, including valuation (Acikalin et al., 2017; Shenhav & Karmarkar, 2019), memory (Euston et al., 2012), and self-referential thought (Mitchell et al., 2005). An individualized frame of reference will enhance the ability of future studies to gauge similarities and differences among brain activity patterns associated with diverse psychological domains.

## Acknowledgements

We thank Lauren DiNicola, Daniel Reznik, Xavier Guell, and David Somers for helpful discussions, and Daniel Sussman for initial guidance on community detection and evaluation methods. This work was supported by grants from the National Science Foundation (BCS-1755757 and BCS-1625552) and the Office of Naval Research (MURI award N00014-16-1-2832 and DURIP award N00014-17-1-2304). Data were provided by the Human Connectome Project, WU-Minn Consortium (Principal Investigators: David Van Essen and Kamil Ugurbil; 1U54MH091657) funded by the 16 NIH Institutes and Centers that support the NIH Blueprint for Neuroscience Research; and by the McDonnell Center for Systems Neuroscience at Washington University.

## Supplemental Materials

**Table 1.**
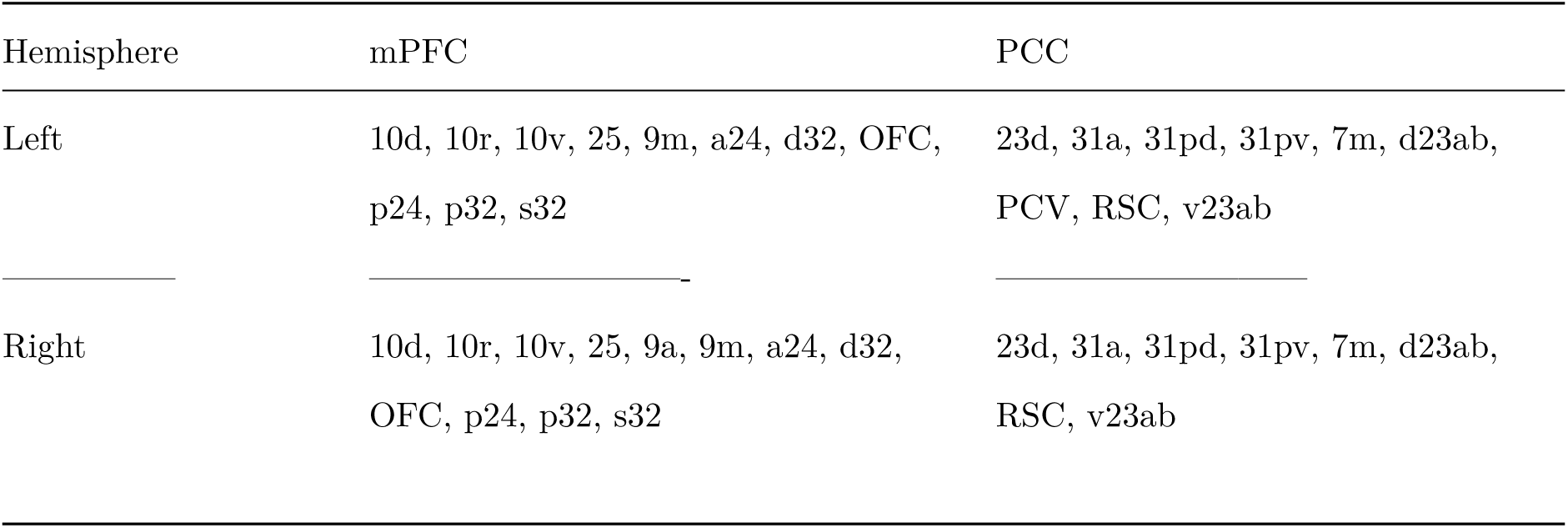
Parcels from Glasser et al. (2016) contained in the search space.

| Hemisphere | mPFC | PCC |
| --- | --- | --- |
| Left | 10d, 10r, 10v, 25, 9m, a24, d32, OFC, p24, p32, s32 | 23d, 31a, 31pd, 31pv, 7m, d23ab, PCV, RSC, v23ab |
| Right | 10d, 10r, 10v, 25, 9a, 9m, a24, d32, OFC, p24, p32, s32 | 23d, 31a, 31pd, 31pv, 7m, d23ab, RSC, v23ab |

**Supplemental Figure 1.**
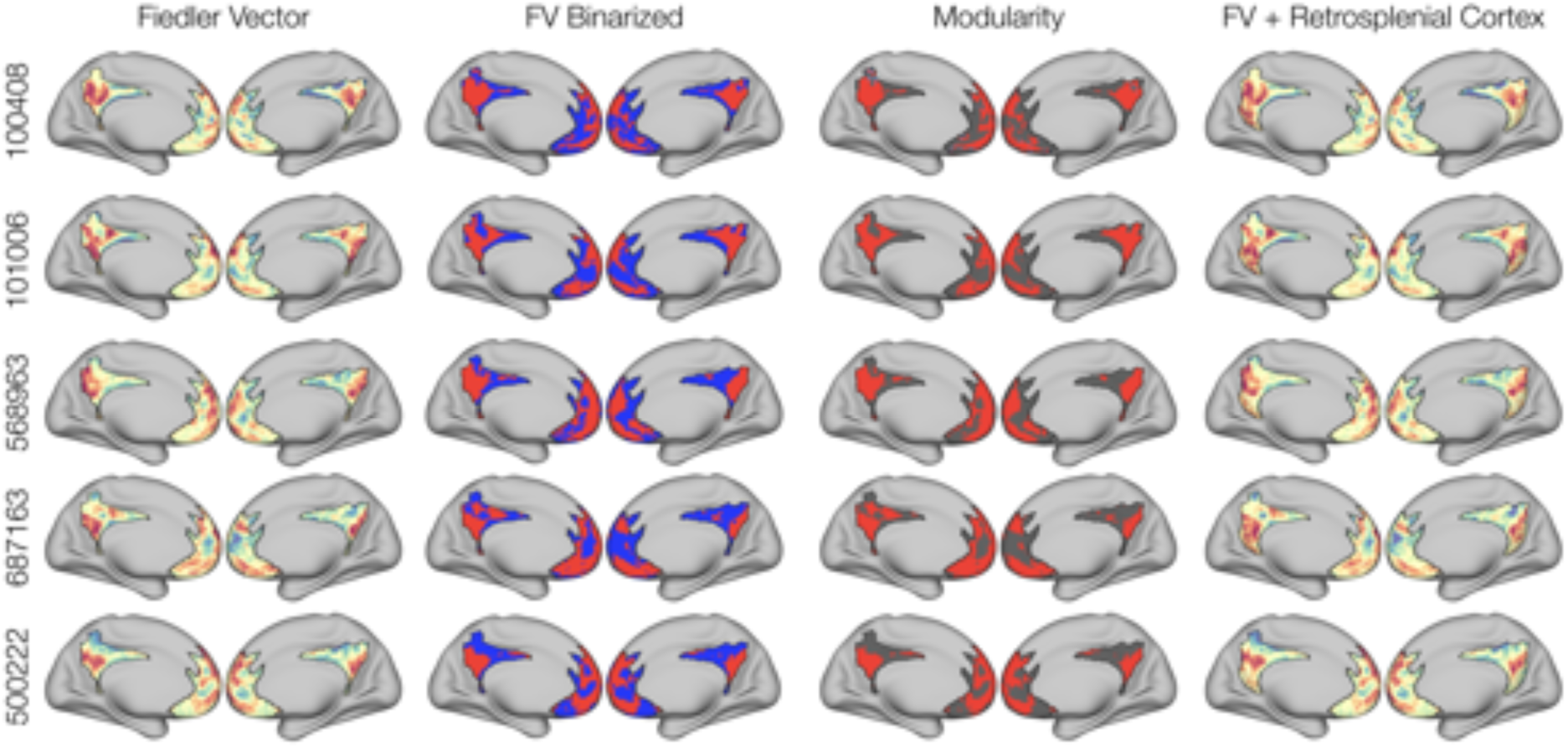
Additional examples of individualized partitionings. The first two columns show both Fiedler vector values and binarized communities, respectively. A common organizational principle is visible, even though it shifts topographically across individuals. The organization is also evident when using Modularity (third column), even though some isolated vertices were sometimes placed in a small third community (see subject 100408). The fourth column shows Fiedler vector maps after adding retrosplenial cortex (a common component of canonical DN affected by task-deactivation). The addition of this area to the search space does not alter the original results, and the retrosplenial region tends to be included in the DN community.

**Supplemental Figure 2.**
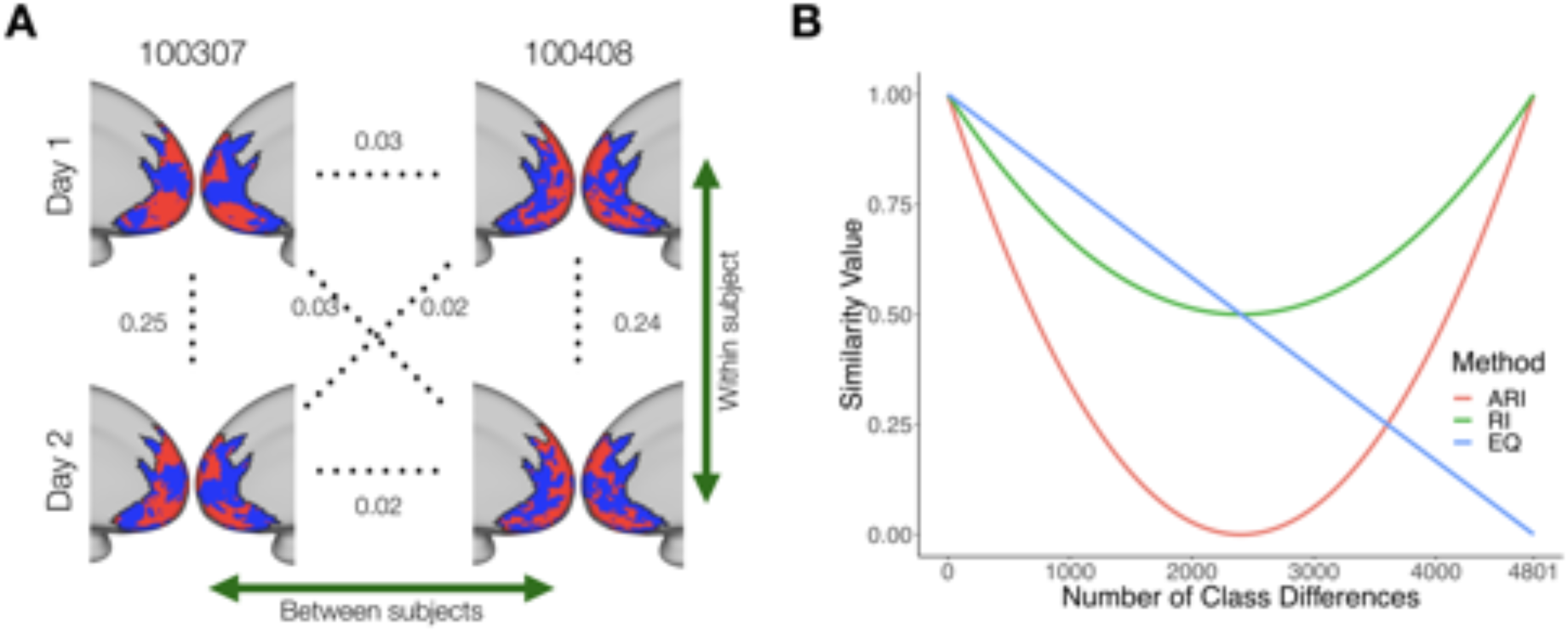
A: Example of an across-day comparison using ARI for two subjects (100307 and 100408). This reflects how qualitatively similar, within-subject partitionings can have relatively small ARI values (here 0.24-0.25), and how partitionings across individuals are much closer to the chance level of zero. B: Simulated comparison between two binary partitionings. The allegiance of each node is progressively switched, and the agreement between the new vector and the original one is computed on each change. The x-axis shows the number of nodes switched. Comparing the increasingly dissimilar maps by computing the proportion of equal cluster labelings (EQ) shows the expected linear decrease in similarity. The unadjusted form of the ARI (RI) displays a nonlinearly decaying similarity, and increases after reaching 50% as a result of node pairs once again being grouped in the same/different clusters (making the index label-agnostic). The ARI decays more steeply as a function of increasing dissimilarity, reaching 0 at chance levels. Low ARI values can therefore still occur when there is systematic agreement between partitionings.} \end{figure}

**Supplemental Figure 3.**
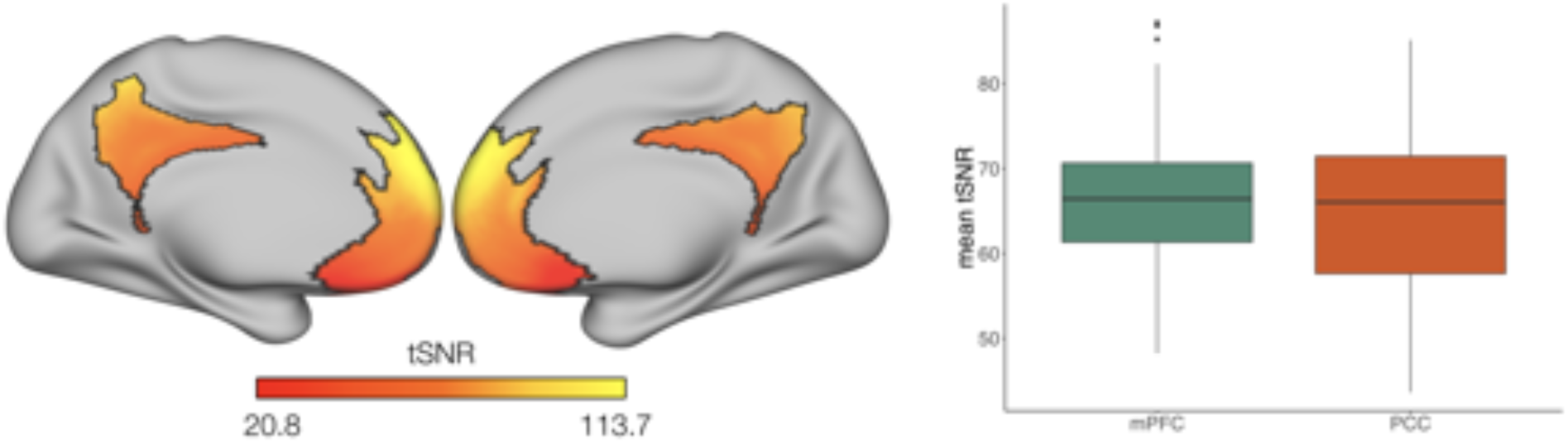
BOLD signal quality in the mPFC and PCC search space. Left: Surface map displaying the vertex-wise mean tSNR across individuals. Right: Mean tSNR for mPFC and PCC across individuals.

**Supplemental Figure 4.**
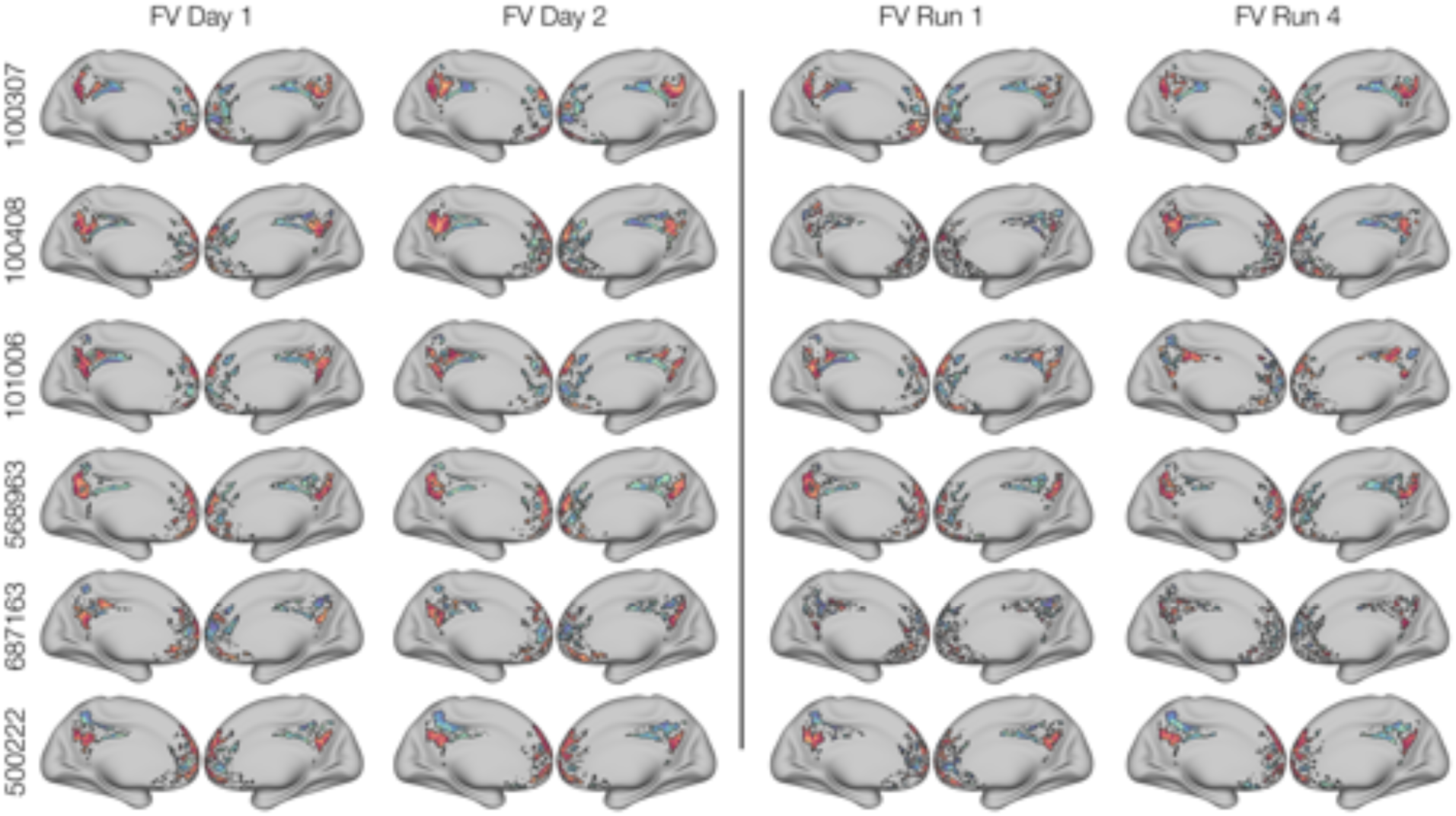
Visual examples of idiosyncratic organization across days and runs. Left: Thresholded partitionings captured with SP were highly similar across days within individuals, but were topographically distinct across individuals. The organization estimated for each day was generally similar to that captured using the full time series (Supplemental Figure 1). Right: Thresholded partitionings estimated using the first scanning run from day 1, and the last scanning run from day 2. As expected, the considerable reduction in the amount of data used decreased the reliability of the community localization across single scanning runs, which were in some cases irrecoverable (e.g. subjects 100408 and 687163). Even so, the general location of DN and non-DN was still observable in many cases, and is comparable to the organization seen using larger amounts of data. This shows that even with notably noisier estimates and lower within-individual ARI values from working with less data, it was possible to gain information about the general location of DN and non-DN in individuals.

**Supplemental Figure 5.**
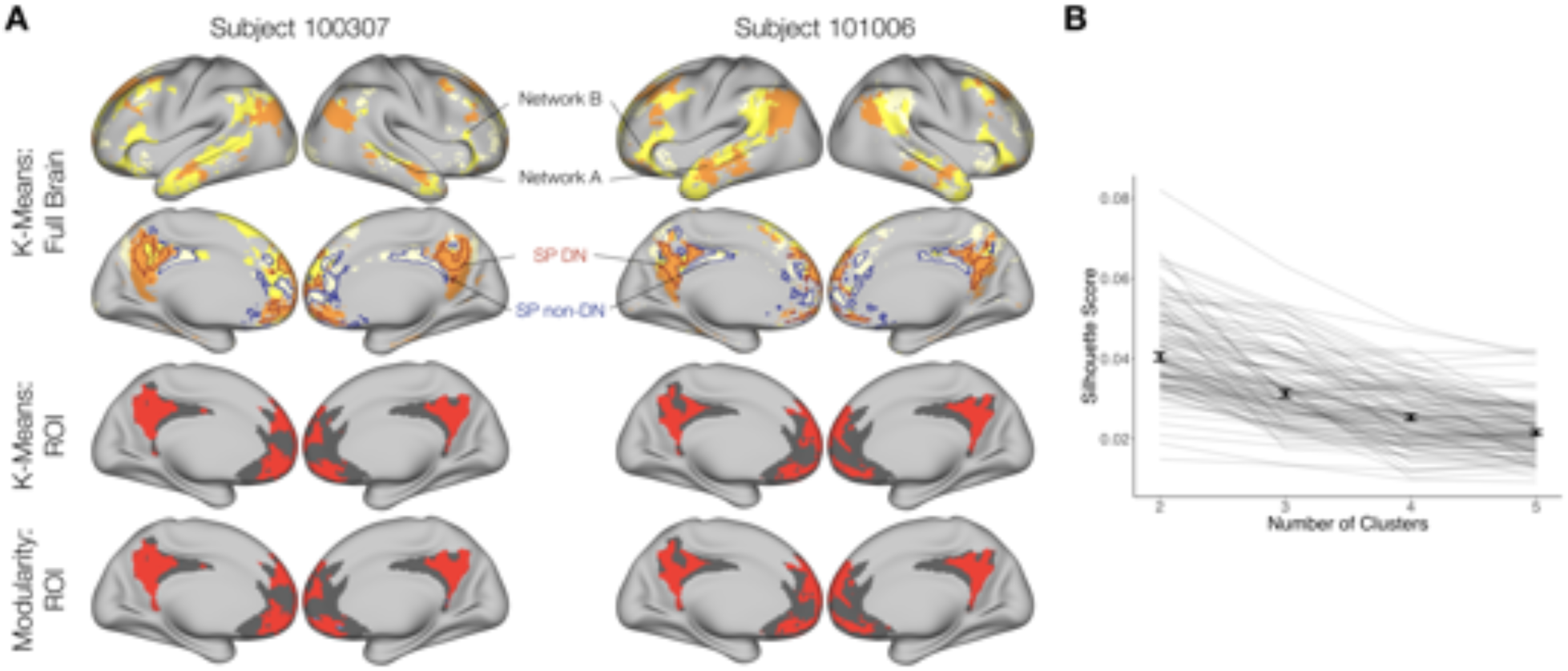
Additional checks to confirm the match between our findings and Braga et al. (2019). A: DN sub-networks A and B identified through whole-brain k-means clustering were congruent with the existence of a third sub-network, which matched the non-DN SP community (top rows). Limiting k-means to two clusters within the search space aligned with results from modularity (bottom two rows). B: K-means based silhouette scores were computed per individual to determine the appropriate number of clusters within the search space (the higher the score, the better). This showed that two networks produced the preferred solution.

## References

Acikalin, M. Y., Gorgolewski, K. J., & Poldrack, R. A. (2017). A coordinate-based meta-analysis of overlaps in regional specialization and functional connectivity across subjective value and default mode networks. Frontiers in Neuroscience, 11 (JAN), 1–11. https://doi.org/10.3389/fnins.2017.00001

Amiez, C., & Petrides, M. (2014). Neuroimaging evidence of the anatomo-functional organization of the human cingulate motor areas. Cerebral Cortex, 24 (3), 563–578. https://doi.org/10.1093/cercor/bhs329

Amiez, C., Neveu, R., Warrot, D., Petrides, M., Knoblauch, K., & Procyk, E. (2013). The location of feedback-related activity in the midcingulate cortex is predicted by local morphology. Journal of Neuroscience, 33 (5), 2217–2228. https://doi.org/10.1523/JNEUROSCI.2779-12.2013

Andrews-Hanna, J. R., Reidler, J. S., Huang, C., & Buckner, R. L. (2010). Evidence for the Default Network’s Role in Spontaneous Cognition. Journal of Neurophysiology, 104 (1), 322–335. https://doi.org/10.1152/jn.00830.2009

Bartra, O., McGuire, J. T., & Kable, J. W. (2013). The valuation system: A coordinate-based meta-analysis of BOLD fMRI experiments examining neural correlates of subjective value. NeuroImage, 76, 412–427. https://doi.org/10.1016/j.neuroimage.2013.02.063

Bassett, D. S., & Sporns, O. (2017). Network neuroscience. Nature Neuroscience, 20 (3), 353–364. https://doi.org/10.1038/nn.4502

Belkin, M., & Niyogi, P. (2003). Laplacian eigenmaps for dimensionalitu reduction and data representation. Neural Computation, 15, 1373–1396. https://doi.org/10.1162/089976603321780317

Braga, R. M., & Buckner, R. L. (2017). Parallel Interdigitated Distributed Networks within the Individual Estimated by Intrinsic Functional Connectivity. Neuron, 95 (2), 457–471.e5. https://doi.org/10.1016/j.neuron.2017.06.038

Braga, R. M., Van Dijk, K. R., Polimeni, J. R., Eldaief, M. C., & Buckner, R. L. (2019). Parallel distributed networks resolved at high resolution reveal close juxtaposition of distinct regions. Journal of Neurophysiology, jn.00808.2018. https://doi.org/10.1152/jn.00808.2018

Buckner, R. L., & Carroll, D. C. (2007). Self-projection and the brain. Trends in Cognitive Sciences, 11 (2), 49–57. https://doi.org/10.1016/j.tics.2006.11.004

Buckner, R. L., & DiNicola, L. M. (2019). The Brain’s Default Network: Updated Anatomy, Physiology, and Evolving Insights. Nature Reviews Neuroscience. https://doi.org/10.1038/s41583-019-0212-7

Buckner, R. L., Andrews-Hanna, J. R., & Schacter, D. L. (2008). The brain’s default network: Anatomy, function, and relevance to disease. Annals of the New York Academy of Sciences, 1124, 1–38. https://doi.org/10.1196/annals.1440.011

Burgess, G. C., Kandala, S., Nolan, D., Laumann, T. O., Power, J. D., Adeyemo, B., … & Barch, D. M. (2016). Evaluation of Denoising Strategies to Address Motion-Correlated Artifacts in Resting-State Functional Magnetic Resonance Imaging Data from the Human Connectome Project. Brain Connectivity, 6 (9), 669–680.

Chung, F. R. K. (1997). Spectral Graph Theory (Vol. 92). American Mathematical Soc.

Clauset, A., Newman, M. E. J., & Moore, C. (2004). Finding community structure in very large networks. 066111, 1–6. https://doi.org/10.1103/PhysRevE.70.066111

Clithero, J. A., & Rangel, A. (2014). Informatic parcellation of the network involved in the computation of subjective value. Social Cognitive and Affective Neuroscience, 9 (9), 1289–1302. https://doi.org/10.1093/scan/nst106

Conroy, B. R., Singer, B. D., Guntupalli, J. S., Ramadge, P. J., & Haxby, J. V. (2013). Inter-subject alignment of human cortical anatomy using functional connectivity. NeuroImage, 81, 400–411. https://doi.org/10.1016/j.neuroimage.2013.05.009

Csardi, G., & Nepusz, T. (2006). The igraph software package for complex network research. InterJournal Complex Systems, 1695.

Dale, A. M., Fischl, B., & Sereno, M. I. (1999). Cortical surface-based analysis: I. Segmentation and surface reconstruction. NeuroImage, 9 (2), 179–194. https://doi.org/10.1006/nimg.1998.0395

De La Vega, A., Chang, L. J., Banich, M. T., Wager, T. D., & Yarkoni, T. (2016). Large-Scale Meta-Analysis of Human Medial Frontal Cortex Reveals Tripartite Functional Organization. The Journal of Neuroscience, 36 (24), 6553–6562. https://doi.org/10.1523/JNEUROSCI.4402-15.2016

DiNicola, L. M., Braga, R. M., & Buckner, R. L. (2019). Parallel Distributed Networks Dissociate Episodic and Social Functions Within the Individual. BioRxiv. https://doi.org/10.1101/733048

Euston, D. R., Gruber, A. J., & McNaughton, B. L. (2012). The role of medial prefrontal cortex in memory and decision making. Neuron, 76 (6), 1057–1070. https://doi.org/10.1016/j.neuron.2012.12.002

Fedorenko, E., Duncan, J., & Kanwisher, N. (2012). Language-selective and domain-general regions lie side by side within Broca’s area. Current Biology, 22 (21), 2059–2062. https://doi.org/10.1016/j.cub.2012.09.011

Fiedler, M. (1975). A Property of Eigenvectors of Nonnegative Symmetric Matrices and its Application to Graph Theory. Czechoslovak Mathematical Journal, 25 (100), 619–633.

Fischl, B., Sereno, M. I., & Dale, A. M. (1999). Cortical surface-based analysis: II. Inflation, flattening, and a surface-based coordinate system. NeuroImage, 9 (2), 195–207. https://doi.org/10.1006/nimg.1998.0396

Fortunato, S., & Hric, D. (2016). Community detection in networks: A user guide. Physics Reports, 659, 1–44. https://doi.org/10.1016/j.physrep.2016.09.002

Fox, M. D., Snyder, A. Z., Vincent, J. L., Corbetta, M., Van Essen, D. C., & Raichle, M. E. (2005). The human brain is intrinsically organized into dynamic, anticorrelated functional networks. Proceedings of the National Academy of Sciences, 102 (27), 9673–9678. https://doi.org/10.1073/pnas.0504136102

Fritsch, A. (2012). mcclust: Process an MCMC Sample of Clusterings. R package version 1.0. Retrieved from https://cran.r-project.org/package=mcclust

Garcia, J. O., Ashourvan, A., Muldoon, S., Vettel, J. M., & Bassett, D. S. (2018). Applications of Community Detection Techniques to Brain Graphs: Algorithmic Considerations and Implications for Neural Function. Proceedings of the IEEE, 106 (5), 846–867. https://doi.org/10.1109/JPROC.2017.2786710

Ghasemian, A., Hosseinmardi, H., & Clauset, A. (2019). Evaluating Overfit and Underfit in Models of Network Community Structure. IEEE Transactions on Knowledge and Data Engineering. https://doi.org/10.1109/TKDE.2019.2911585

Gkantsidis, C., Mihail, M., & Zegura, E. (2003). Spectral analysis of Internet topologies. Proc. IEEE INFOCOM, 00 (C), 364–374. https://doi.org/10.1109/INFCOM.2003.1208688

Glasser, M. F., Coalson, T. S., Robinson, E. C., Hacker, C. D., Harwell, J., Yacoub, E., … Van Essen, D. C. (2016). A multi-modal parcellation of human cerebral cortex. Nature, 536 (7615), 171–178. https://doi.org/10.1038/nature18933

Glasser, M. F., Sotiropoulos, S. N., Wilson, J. A., Coalson, T. S., Fischl, B., Andersson, J. L., … Jenkinson, M. (2013). The minimal preprocessing pipelines for the Human Connectome Project. NeuroImage, 80, 105–124. https://doi.org/10.1016/j.neuroimage.2013.04.127

Gordon, E. M., Laumann, T. O., Gilmore, A. W., Newbold, D. J., Greene, D. J., Berg, J. J., … Dosenbach, N. U. (2017). Precision Functional Mapping of Individual Human Brains. Neuron, 95 (4), 791–807.e7. https://doi.org/10.1016/j.neuron.2017.07.011

Gratton, C., Laumann, T. O., Nielsen, A. N., Greene, D. J., Gordon, E. M., Gilmore, A. W., … Petersen, S. E. (2018). Functional Brain Networks Are Dominated by Stable Group and Individual Factors, Not Cognitive or Daily Variation. Neuron, 439–452. https://doi.org/10.1016/j.neuron.2018.03.035

Greicius, M. D., Krasnow, B., Reiss, A. L., & Menon, V. (2003). Functional connectivity in the resting brain: A network analysis of the default mode hypothesis. 100 (1), 253–258. https://doi.org/10.1073/pnas.0135058100

Griffanti, L., Salimi-Khorshidi, G., Beckmann, C. F., Auerbach, E. J., Douaud, G., Sexton, C. E., … Smith, S. M. (2014). ICA-based artefact removal and accelerated fMRI acquisition for improved resting state network imaging. NeuroImage, 95, 232–247. https://doi.org/10.1016/j.neuroimage.2014.03.034

Guntupalli, J. S., Feilong, M., & Haxby, J. V. (2018). A computational model of shared fine-scale structure in the human connectome. PLoS Computational Biology, 14 (4), 1–26. https://doi.org/10.1371/journal.pcbi.1006120

Gusnard, D. A., Akbudak, E., Shulman, G. L., & Raichle, M. E. (2001). Medial prefrontal cortex and self-referential mental activity: Relation to a default mode of brain function. Proceedings of the National Academy of Sciences, 98 (7), 4259–4264. https://doi.org/10.1073/pnas.071043098

Hassabis, D., & Maguire, E. A. (2007). Deconstructing episodic memory with construction. Trends in Cognitive Sciences, 11 (7), 299–306. https://doi.org/10.1016/j.tics.2007.05.001

Higham, D. J., Kalna, G., & Kibble, M. (2007). Spectral clustering and its use in bioinformatics. Journal of Computational and Applied Mathematics, 204 (1), 25–37. https://doi.org/10.1016/j.cam.2006.04.026

Hiser, J., & Koenigs, M. (2018). The Multifaceted Role of the Ventromedial Prefrontal Cortex in Emotion, Decision Making, Social Cognition, and Psychopathology. Biological Psychiatry, 83 (8), 638–647. https://doi.org/10.1016/j.biopsych.2017.10.030

Hubert, L., & Arabie, P. (1985). Comparing partitions. Journal of Classification, 2 (1), 193–218. https://doi.org/10.1007/BF01908075

Huntenburg, J. M., Bazin, P. L., & Margulies, D. S. (2018). Large-Scale Gradients in Human Cortical Organization. Trends in Cognitive Sciences, 22 (1), 21–31. https://doi.org/10.1016/j.tics.2017.11.002

Kable, J. W., & Glimcher, P. W. (2007). The neural correlates of subjective value during intertemporal choice. Nature Neuroscience, 10 (12), 1625–1633.

Kernbach, J. M., Yeo, B. T. T., Smallwood, J., Margulies, D. S., Thiebaut de Schotten, M., Walter, H., … Bzdok, D. (2018). Subspecialization within default mode nodes characterized in 10,000 UK Biobank participants. Proceedings of the National Academy of Sciences, (November), 201804876. https://doi.org/10.1073/pnas.1804876115

Kong, R., Li, J., Orban, C., Sabuncu, M. R., Liu, H., Schaefer, A., … Yeo, B. T. T. (2018). Spatial Topography of Individual-Specific Cortical Networks Predicts Human Cognition, Personality, and Emotion. Cerebral Cortex, (May 2018), 2533–2551. https://doi.org/10.1093/cercor/bhy123

Kragel, P. A., Kano, M., Van Oudenhove, L., Ly, H. G., Dupont, P., Rubio, A., … Wager, T. D. (2018). Generalizable representations of pain, cognitive control, and negative emotion in medial frontal cortex. Nature Neuroscience, 1. https://doi.org/10.1038/s41593-017-0051-7

Laird, A. R., Eickhoff, S. B., Li, K., Robin, D. A., Glahn, D. C., & Fox, P. T. (2009). Investigating the Functional Heterogeneity of the Default Mode Network Using Coordinate-Based Meta-Analytic Modeling. Journal of Neuroscience, 29 (46), 14496–14505. https://doi.org/10.1523/JNEUROSCI.4004-09.2009

Laumann, T. O., Gordon, E. M., Adeyemo, B., Snyder, A. Z., Joo, S. J., Chen, M. Y., … Petersen, S. E. (2015). Functional System and Areal Organization of a Highly Sampled Individual Human Brain. Neuron, 87 (3), 658–671. https://doi.org/10.1016/j.neuron.2015.06.037

Levy, I., Lazzaro, S. C., Rutledge, R. B., & Glimcher, P. W. (2011). Choice from Non-Choice: Predicting Consumer Preferences from Blood Oxygenation Level-Dependent Signals Obtained during Passive Viewing. Journal of Neuroscience, 31 (1), 118–125. https://doi.org/10.1523/JNEUROSCI.3214-10.2011

Lopez-Persem, A., Verhagen, L., Amiez, C., Petrides, M., & Sallet, J. (2019). The human ventromedial prefrontal cortex: sulcal morphology and its influence on functional organization. The Journal of Neuroscience, 39 (19), 2060–2018. https://doi.org/10.1523/JNEUROSCI.2060-18.2019

Mackey, S., & Petrides, M. (2014). Architecture and morphology of the human ventromedial prefrontal cortex. European Journal of Neuroscience, 40 (5), 2777–2796. https://doi.org/10.1111/ejn.12654

Margulies, D. S., Ghosh, S. S., Goulas, A., Falkiewicz, M., Huntenburg, J. M., Langs, G., … Smallwood, J. (2016). Situating the default-mode network along a principal gradient of macroscale cortical organization. Proceedings of the National Academy of Sciences, 113 (44), 12574–12579. https://doi.org/10.1073/pnas.1608282113

Mars, R. B., Passingham, R. E., & Jbabdi, S. (2018). Connectivity Fingerprints: From Areal Descriptions to Abstract Spaces. Trends in Cognitive Sciences, 22 (11), 1026–1037. https://doi.org/10.1016/j.tics.2018.08.009

McKiernan, K. A., Kaufman, J. N., Kucera-Thompson, J., & Binder, J. R. (2003). A Parametric Manipulation of Factors Affecting Task-induced Deactivation in Functional Neuroimaging. Journal of Cognitive Neuroscience, 15, 394–408. https://doi.org/10.1162/089892903321593117

Michalka, S. W., Kong, L., Rosen, M. L., Shinn-Cunningham, B. G., & Somers, D. C. (2015). Short-Term Memory for Space and Time Flexibly Recruit Complementary Sensory-Biased Frontal Lobe Attention Networks. Neuron, 87 (4), 882–892. https://doi.org/10.1016/j.neuron.2015.07.028

Mitchell, J. P., Banaji, M. R., & Macrae, C. N. (2005). The link between social cognition and self-referential thought in the medial prefrontal cortex. Journal of Cognitive Neuroscience, 17 (8), 1306–1315.

Mueller, S., Wang, D., Fox, M. D., Yeo, B. T. T., Sepulcre, J., Sabuncu, M. R., … Liu, H. (2013). Individual Variability in Functional Connectivity Architecture of the Human Brain. Neuron, 77 (3), 586–595. https://doi.org/10.1016/j.neuron.2012.12.028

Nenning, K. H., Liu, H., Ghosh, S. S., Sabuncu, M. R., Schwartz, E., & Langs, G. (2017). Diffeomorphic functional brain surface alignment: Functional demons. NeuroImage, 156 (December 2016), 456–465. https://doi.org/10.1016/j.neuroimage.2017.04.028

Northoff, G., & Hayes, D. J. (2011). Is our self nothing but reward? Biological Psychiatry, 69, 1019–1025. https://doi.org/10.1016/j.biopsych.2010.12.014

Osher, D. E., Saxe, R. R., Koldewyn, K., Gabrieli, J. D., Kanwisher, N., & Saygin, Z. M. (2016). Structural Connectivity Fingerprints Predict Cortical Selectivity for Multiple Visual Categories across Cortex. Cerebral Cortex, 26 (4), 1668–1683. https://doi.org/10.1093/cercor/bhu303

Passingham, R. E., Stephan, K. E., & Kötter, R. (2002). The anatomical basis of functional localization in the cortex. Nature Reviews Neuroscience, 3 (8), 606–616. https://doi.org/10.1038/nrn893

Power, J. D., Mitra, A., Laumann, T. O., Snyder, A. Z., Schlaggar, B. L., & Petersen, S. E. (2014). Methods to detect, characterize, and remove motion artifact in resting state fMRI. NeuroImage, 84, 320–341. https://doi.org/10.1016/j.neuroimage.2013.08.048

R Core Computing Team. (2017). R: A Language and Environment for Statistical Computing. Retrieved from https://www.r-project.org/

Reddan, M. C., Wager, T. D., & Schiller, D. (2018). Attenuating Neural Threat Expression with Imagination. Neuron, 100 (4), 994–1005.e4. https://doi.org/10.1016/j.neuron.2018.10.047

Rubinov, M., & Sporns, O. (2010). Complex network measures of brain connectivity: Uses and interpretations. NeuroImage, 52 (3), 1059–1069. https://doi.org/10.1016/j.neuroimage.2009.10.003

Salimi-Khorshidi, G., Douaud, G., Beckmann, C. F., Glasser, M. F., Griffanti, L., & Smith, S. M. (2014). Automatic denoising of functional MRI data: Combining independent component analysis and hierarchical fusion of classifiers. NeuroImage, 90, 449–468. https://doi.org/10.1016/j.neuroimage.2013.11.046

Saygin, Z. M., Osher, D. E., Koldewyn, K., Reynolds, G., Gabrieli, J. D., & Saxe, R. R. (2011). Anatomical connectivity patterns predict face selectivity in the fusiform gyrus. Nature Neuroscience, 15 (2), 321–327. https://doi.org/10.1038/nn.3001

Saygin, Z. M., Osher, D. E., Norton, E. S., Youssoufian, D. A., Beach, S. D., Feather, J., … Kanwisher, N. (2016). Connectivity precedes function in the development of the visual word form area. Nature Neuroscience, 19 (9), 1250–1255. https://doi.org/10.1038/nn.4354

Schacter, D. L., Addis, D. R., & Buckner, R. L. (2007). Remembering the past to imagine the future: The prospective brain. Nature Reviews Neuroscience, 8 (9), 657–661. https://doi.org/10.1038/nrn2213

Schiller, D., Levy, I., Niv, Y., LeDoux, J. E., & Phelps, E. A. (2008). From fear to safety and back: Reversal of fear in the human brain. Journal of Neuroscience, 28 (45), 11517–11525. https://doi.org/10.1523/JNEUROSCI.2265-08.2008

Shenhav, A., & Karmarkar, U. R. (2019). Dissociable components of the reward circuit are involved in appraisal versus choice. Scientific Reports, 9 (1), 172320. https://doi.org/10.1038/s41598-019-38927-7

Smith, S. M., Fox, P. T., Miller, K. L., Glahn, D. C., Fox, P. M., Mackay, C. E., … Beckmann, C. F. (2009). Correspondence of the brain’s functional architecture during activation and rest. Proceedings of the National Academy of Sciences, 106 (31), 13040 LP–13045. https://doi.org/10.1073/pnas.0905267106

Spreng, N. (2012). The fallacy of a “task-negative” network. Frontiers in Psychology, 3 (MAY), 1–5. https://doi.org/10.3389/fpsyg.2012.00145

Tian, Y., & Zalesky, A. (2018). Characterizing the functional connectivity diversity of the insula cortex: Subregions, diversity curves and behavior. NeuroImage, 183, 716–733. https://doi.org/10.1016/j.neuroimage.2018.08.055

Tobyne, S. M., Osher, D. E., Michalka, S. W., & Somers, D. C. (2017). Sensory-biased attention networks in human lateral frontal cortex revealed by intrinsic functional connectivity. NeuroImage, 162 (August), 362–372. https://doi.org/10.1016/j.neuroimage.2017.08.020

Tobyne, S. M., Somers, D. C., Brissenden, J. A., Michalka, S. W., Noyce, A. L., & Osher, D. E. (2018). Prediction of individualized task activation in sensory modality-selective frontal cortex with ‘connectome fingerprinting’. NeuroImage, 183 (August), 173–185. https://doi.org/10.1016/j.neuroimage.2018.08.007

Toker, D., & Sommer, F. T. (2019). Information Integration In Large Brain Networks. PLoS Computational Biology, 15 (2). https://doi.org/10.1371/journal.pcbi.1006807

Van Essen, D. C., Ugurbil, K., Auerbach, E., Barch, D., Behrens, T. E., Bucholz, R., … Yacoub, E. (2012). The Human Connectome Project: A data acquisition perspective. NeuroImage, 62 (4), 2222–2231. https://doi.org/10.1016/j.neuroimage.2012.02.018

Wager, T. D., Lindquist, M. A., Nichols, T. E., Kober, H., & Van Snellenberg, J. X. (2009). Evaluating the consistency and specificity of neuroimaging data using meta-analysis. NeuroImage, 45 (1 Suppl), S210–S221. https://doi.org/10.1016/j.neuroimage.2008.10.061

Wang, D., Buckner, R. L., Fox, M. D., Holt, D. J., Holmes, A. J., Stoecklein, S., … Liu, H. (2015). Parcellating cortical functional networks in individuals. Nature Neuroscience, 18 (12), 1853–1860. https://doi.org/10.1038/nn.4164

Woo, C.-W., Koban, L., Kross, E., Lindquist, M. A., Banich, M. T., Ruzic, L., … Wager, T. D. (2014). Separate neural representations for physical pain and social rejection. Nature Communications, 5 (May), 1–12. https://doi.org/10.1038/ncomms6380

Yeo, B. T. T., Krienen, F. M., Sepulcre, J., Sabuncu, M. R., Lashkari, D., Hollinshead, M., … Buckner, R. L. (2011). The organization of the human cerebral cortex estimated by intrinsic functional connectivity. Journal of Neurophysiology, 106, 1125–1165. https://doi.org/10.1152/jn.00338.2011

Zilles, K., Palomero-Gallagher, N., & Amunts, K. (2013). Development of cortical folding during evolution and ontogeny. Trends in Neurosciences, 36 (5), 275–284. https://doi.org/10.1016/j.tins.2013.01.006

Zlatkina, V., Amiez, C., & Petrides, M. (2016). The postcentral sulcal complex and the transverse postcentral sulcus and their relation to sensorimotor functional organization. European Journal of Neuroscience, 43 (10), 1268–1283. https://doi.org/10.1111/ejn.13049

